# A tRNA-derived small RNA orchestrates cardioprotection through distinct cell-type-specific mechanisms

**DOI:** 10.64898/2026.09.28.755177

**Authors:** Tian Hao, Jin Li, Kaushik V. Amancherla, Lingfei Sun, Priyanka Gokulnath, Alejandra Patino-Guerrero, Ritin Sharma, Krystine Garcia-Mansfield, Chunyang Xiao, Aidan C. Manning, Emeli Chartterjee, Xiaoting Yu, Richard Xu, Yiming Zhang, Zhihong Yang, Pavel Ivanov, John J. Rossi, William A. Goddard, Hua Zhu, Oluwaseun Akeju, Eric R. Gamazon, Wuqiang Zhu, Patrick Pirrotte, Ravi Shah, Junjie Xiao, Saumya Das, Guoping Li

## Abstract

Pathological cardiac fibrosis and hypertrophy, key contributors to adverse remodeling that drives heart failure (HF), are not adequately targeted by current therapies. We identified tRNA-Asp-GTC-3′tDR, an ischemia-induced tRNA-derived small RNA (tDR), as an endogenous regulator of both remodeling processes across cardiac fibroblasts and cardiomyocytes. In murine models of myocardial ischemia- and pressure overload-induced HF, cardiac delivery of tRNA-Asp-GTC-3′tDR reduced fibrosis and hypertrophy, preserved cardiac function, and improved survival, with limited three doses conferring durable protection after ischemia, whereas silencing it worsened both. Strikingly, tRNA-Asp-GTC-3’tDR acts through distinct mechanisms in different cell types: in fibroblasts, it assembles stress granules that sequester ribosome-loaded profibrotic transcripts, whereas in cardiomyocytes it engages pseudouridine synthase PUS7 to activate RNA autophagy. Its regulation and functions were confirmed in human cardiac cells and tissues. These findings demonstrate that a single tDR coordinates cell-type-specific protective programs in the heart, suggesting a new therapeutic axis to mitigate HF progression.

## INTRODUCTION

The global prevalence of heart failure (HF) is estimated to be 55-64 million patients, or 1-3% of the population. HF continues to have a significant impact, contributing to 425,147 deaths in the U.S. in 2022 (45% of all cardiovascular deaths), with mortality rising since 2012 ^1–3^. While wider adoption of guideline-directed medical therapy (GDMT) has improved clinical outcomes in chronic HF patients ^4,5^, patients on GDMT still carry substantial residual risk of HF progression ^6,7^, indicating that key molecular drivers of HF pathogenesis are not directly addressed by current GDMT.

The stress signals that precipitate the tissue remodeling underlying HF progression act through different pathways in different cardiac cell types. Two remodeling processes that dominate this response, yet are not specifically targeted by GDMT, are the conversion of cardiac fibroblasts (CFs) into matrix-secreting myofibroblasts and pathological cardiomyocyte (CM) hypertrophy^8,9^. These responses, executed by distinct molecular programs in these two cell types, advance in parallel and together drive the structural and functional deterioration that culminates in overt HF and its complications^10^. Myocardial fibrosis stiffens the myocardium, provides a substrate for ventricular arrhythmias, and impairs cardiac function, predicting adverse outcomes in patients ^11–13^, while pathological CM hypertrophy contributes comparably to contractile dysfunction and disease progression^14^. Consistent with their pathogenic roles, experimental approaches that limit maladaptive fibrosis^15,16^ or hypertrophy^17^ mitigate adverse cardiac remodeling and HF progression. These observations suggest that identifying and targeting novel regulatory molecules of these complementary remodeling pathways could meaningfully reduce the residual risk that persists in HF patients.

Transfer RNA-derived small RNAs (tsRNAs or tDRs) constitute an abundant and evolutionarily conserved class of RNAs, which are generated by the ribonuclease-mediated cleavages of tRNAs in a manner shaped by both cellular state and tRNA tertiary structure and modifications ^18,19^. Rather than inert degradation products, tDRs are functional regulatory RNAs that reshape translation, RNA metabolism, and cellular adaptation to stress, with established roles in tumor progression and neurological disorders ^20,21^. tDRs are abundantly expressed in cardiac cells and tissues and can also be released into the circulation ^22^, where they have been explored primarily as biomarkers of cardiovascular stress ^23^, while their endogenous regulation and functional roles in cardiovascular disease are only beginning to be explored.

Under stress conditions, cells typically produce tDRs as an adaptive mechanism ^24,25^, and two of their activities have emerged as protective in disease. The first is the assembly of stress granules (SGs), cytoplasmic ribonucleoprotein condensates that sequester non-translating mRNAs and reprogram protein synthesis during stress ^26^. Terminal oligo-guanine motif-containing tDRs, including the 5′ halves of tRNA-Ala-AGC and tRNA-Cys-GCA, fold into G-quadruplex (G4) structures and drive SG formation ^27,28^. In motor neurons, this activity has been linked to protection against neurodegeneration ^29^. The second is RNA autophagy. We recently found that hypoxia-induced Asp-GTC-3’tDR, derived from the 3’ end of tRNA-Asp-GTC, sequesters pseudouridine synthase 7 (PUS7) from its histone mRNA substrates and routes these pseudouridine-deficient transcripts to autophagosomes for degradation, triggering RNA autophagy that protects kidney cells from injury ^30^. These examples establish that stress-induced tDRs can direct distinct regulatory pathways to enhance cellular adaptation, yet their roles in the heart have not yet been examined.

Here, through a screen of ischemia-regulated tDRs for cardioprotective activity, we identify Asp-GTC-3′tDR as an endogenous regulator of distinct cardioprotective pathways in preclinical models of HF. Asp-GTC-3′tDR is acutely induced by ischemic stress in both CFs and CMs, but its subsequent decline in expression coincides with myofibroblast activation and CM hypertrophy. In cultured cardiac cells, Asp-GTC-3’tDR exerts robust anti-fibrotic and anti-hypertrophic effects. Cardiac delivery of Asp-GTC-3’tDR suppresses both pathological fibrosis and hypertrophy, preserves contractile function, and improves survival in models of myocardial infarction (MI), ischemia/reperfusion (I/R), and pressure overload; conversely, depleting it provokes both maladaptive programs in the healthy heart. Mechanistically, the same tDR orchestrates distinct molecular signaling in the two primary cardiac cell types: in CFs, it nucleates SGs that selectively sequester profibrotic transcripts and restrain myofibroblast activation, revealing an unanticipated antifibrotic function for SGs in the heart; in CMs, it sequesters PUS7 to prevent pseudouridylation of histone mRNAs, enhancing RNA autophagy and thereby limiting hypertrophic growth. These mechanisms are preserved in human cardiac cells, and Asp-GTC-3′tDR is elevated in the myocardium of patients with ischemic cardiomyopathy (ICM). Our findings establish that a single stress-induced tDR can deploy distinct RNA-regulatory machineries in different cells of the same organ to produce a unified protective outcome, positioning Asp-GTC-3′tDR as a therapeutic candidate that addresses key remodeling pathways in HF and may complement current GDMT.

## RESULTS

### Ischemia-induced Asp-GTC-3′tDR attenuates myofibroblast activation in cultured CFs

Cardiac fibrosis, driven by the activation of resident fibroblasts into matrix-secreting myofibroblasts, is a central contributor to adverse remodeling and HF^31^. Although canonical pathways of myofibroblast activation, such as the TGF-β/SMAD pathway, have been characterized in detail, emerging work points to complementary regulatory axes ^32,33^. Ischemic myocardial injury elicits an inflammatory and TGF-β-rich response that drives myofibroblast activation ^34^. Previously, we identified ischemia as a potent regulator of tDR biogenesis ^22^. We therefore hypothesized that ischemia-regulated tDRs might be important modulators of myofibroblast activation. To identify the tDR candidates most likely to be causally linked to the fibrotic response to ischemic stress, we leveraged our stress-specific cellular tDR atlas to identify the tDRs most significantly induced by ischemia in CFs^22^. AlkB-facilitated RNA methylation sequencing (ARM-seq) had identified 24 significantly upregulated tDRs in CFs after 5 hours of ischemia (**Figure 1A, Table S1**). We validated the 5 most highly upregulated tDRs, including Glu-TTC-5’tDR (tDR-1:32-Glu-TTC-4), Glu-CTC-5’tDR (tDR-1:32-Glu-CTC-1), Asp-GTC-5’tDR (tDR-1:32-Asp-GTC-2-M2), Asp-GTC-3’tDR (tDR-39:72-Asp-GTC-2), and Gly-GCC-5’tDR (tDR-1:32-Gly-GCC-2-M2), using Northern blot analysis (**Figure 1B**). To determine which of these tDRs may regulate myofibroblast activation, we transfected primary neonatal rat CFs (NRCFs) with each synthetic tDR mimic and measured the expression of profibrotic genes. Among the five, Asp-GTC-3’tDR potently suppressed profibrotic gene expression (**Figure 1C**), suggesting it acts as an endogenous ‘brake’ on myofibroblast activation. We therefore focused on it for further functional and mechanistic studies.

**Figure 1.**
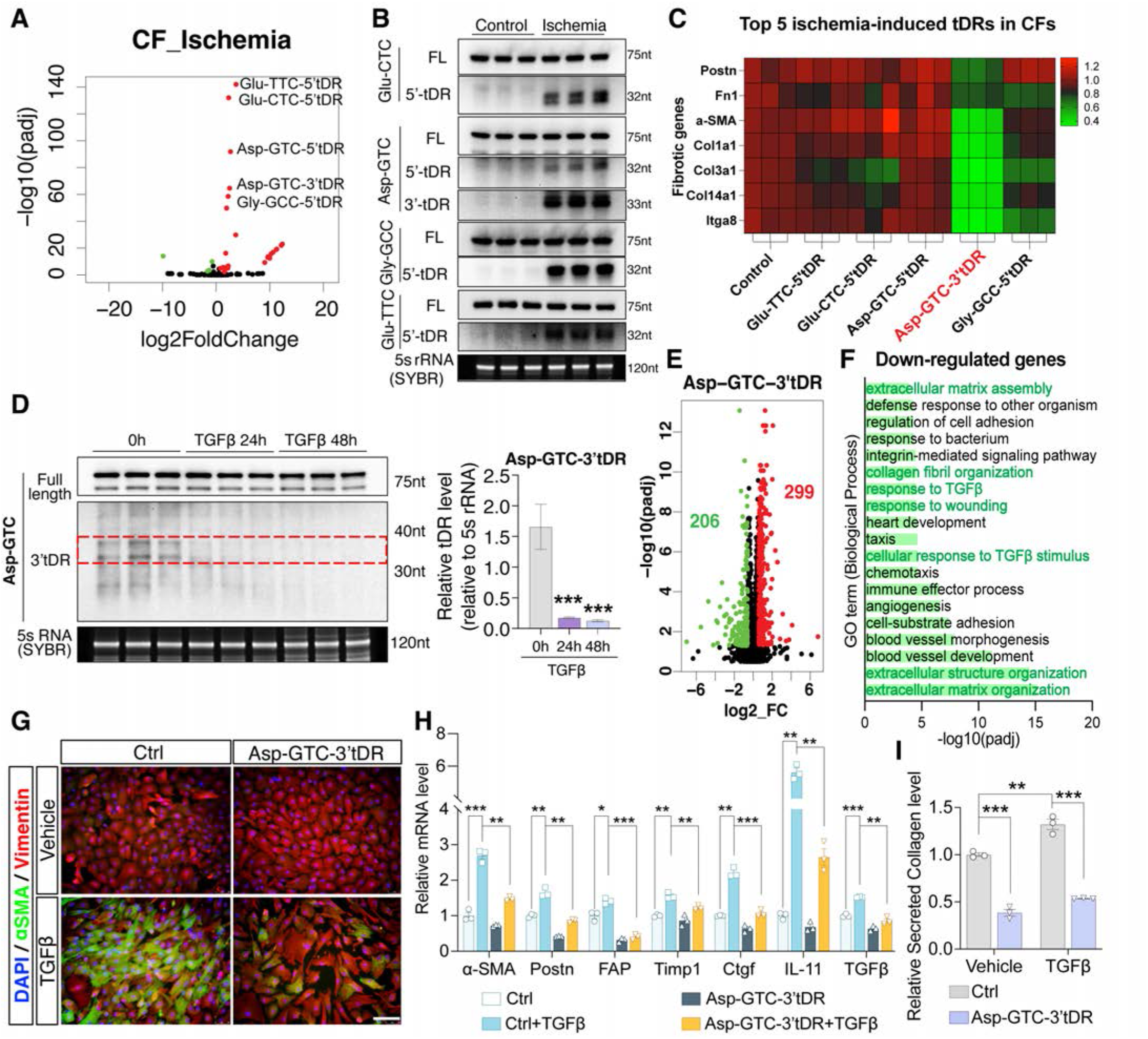
Ischemia-induced Asp-GTC-3′tDR attenuates myofibroblast activation in cultured CFs. **(A)** Volcano plot of differentially expressed tDRs in NRCFs under ischemia versus untreated controls. The top 5 upregulated tDRs are indicated. **(B)** Northern blot analysis of the top 5 upregulated tDRs in NRCFs following 5 hours of ischemia, showing full-length (FL) tRNAs and their corresponding 5’tDRs and/or 3’tDRs. 5S rRNA stained with SYBR Gold served as the loading control. **(C)** Heatmap showing the effects of the top 5 ischemia-induced tDRs on profibrotic gene expression following transfection of their synthetic mimics into NRCFs. A mimic with a scrambled sequence was used as control (Ctrl). **(D)** Northern blot analysis and quantification of Asp-GTC-3′tDR levels in NRCFs following TGFβ stimulation. **(E)** Volcano plot of differentially expressed genes in NRCFs transfected with Asp-GTC-3′tDR mimics versus Ctrl mimics. **(F)** Gene Ontology analysis of 206 Asp-GTC-3’tDR-downregulated genes, showing enrichment in extracellular matrix organization/assembly and TGFβ signaling pathways. **(G)** Immunofluorescence staining of NRCFs transfected with Ctrl or Asp-GTC-3′tDR mimic and treated with vehicle or TGFβ for 48 hours. Cells were stained for α-SMA (green), Vimentin (red), and nuclei (DAPI, blue). Scale bar, 100 μm. **(H)** qPCR analysis of myofibroblast activation-associated gene expression in NRCFs transfected with Ctrl or Asp-GTC-3′tDR mimic and treated with vehicle or TGFβ for 48 hours. **(I)** Secreted collagen levels in culture supernatants of NRCFs transfected with Ctrl or Asp-GTC-3’tDR mimics and treated with vehicle or TGFβ for 48 hours. Data are presented as mean ± SEM. Statistical significance was determined by the unpaired two-tailed Student’s t-test. *\*P* < 0.05, *\*\* P*< 0.01, *\*\*\*P* < 0.001.

To distinguish stress-induced tRNA cleavage from de novo transcription, we used a self-quenched Asp-GTC-3′tDR biogenesis reporter^35^, in which a quencher (BHQ) and a fluorophore (FITC) were conjugated to the 5’ and 3’ ends of synthetic full-length tRNA-Asp-GTC, so that fluorescence is released only when the tRNA is cleaved and the resulting fragments separate (**Figure S1A**). Live-cell imaging of reporter-transfected NRCFs revealed progressive fluorescence during ischemic stress, with detectable fluorescence appearing after approximately 75–80 minutes of ischemia (**Figure S1B**), and 5 hours of ischemia significantly increased fluorescence signals relative to normal controls (**Figure S1C**). These data confirmed that ischemic stress drives cleavage of tRNA-Asp-GTC to generate Asp-GTC-3′tDR. The 5’ end of Asp-GTC-3’tDR mapped to position 37, within the anti-codon loop, and Angiogenin (ANG) is a ribonuclease known to cleave tRNAs at the anti-codon loop ^36^. To test the role of ANG in Asp-GTC-3’tDR biogenesis, we knocked down ANG in NRCFs with siRNAs and exposed the cells to ischemic stress (**Figure S1D**). Northern blot analysis showed that ANG silencing substantially impaired ischemia-induced Asp-GTC-3’tDR generation (**Figure S1E**), identifying ANG as a critical ribonuclease for Asp-GTC-3’tDR biogenesis in CFs under ischemia.

We next determined the functional role of Asp-GTC-3′tDR in myofibroblast activation. As CFs activated toward myofibroblasts under TGFβ stimulation, Northern blot showed that Asp-GTC-3′tDR generation was progressively depleted (**Figure 1D**), suggesting that the loss of Asp-GTC-3′tDR expression accompanies profibrotic activation and that this tDR may therefore function as an endogenous suppressor of fibrosis. To characterize the molecular pathways regulated by this tDR, we overexpressed Asp-GTC-3’tDR in NRCFs by transfecting cells with synthetic Asp-GTC-3’tDR and control mimics and profiled the resulting transcriptomic changes. We identified 299 upregulated and 206 downregulated genes in the Asp-GTC-3’tDR group (**Figure 1E**). Gene Ontology analysis linked the downregulated genes to TGFβ signaling and extracellular matrix/structural organization pathways (**Figure 1F**), two functional hallmarks of cardiac fibrosis^37^, consistent with an antifibrotic function. We tested this directly in the TGFβ-induced myofibroblast activation model. NRCFs were transfected with Asp-GTC-3’tDR or control mimics and then stimulated with 10 ng/mL TGFβ for 48 hours. Asp-GTC-3′tDR overexpression suppressed the TGFβ-induced differentiation of CFs into α-SMA-positive myofibroblasts (**Figure 1G**), attenuated the TGFβ-driven induction of fibrotic genes (**Figure 1H**), and reduced extracellular collagen production under both basal and profibrotic conditions (**Figure 1I**). The same antifibrotic effects of Asp-GTC-3’tDR were observed in cultured adult rat CFs under TGFβ treatment, including the inhibited differentiation to α-SMA-positive myofibroblasts, attenuated fibrotic gene expression, and reduced collagen secretion (**Figures S2A-S2C**). Together, these results demonstrated that ischemia-induced Asp-GTC-3’tDR attenuates cardiac fibrosis in cultured CFs.

### Cardiac delivery of Asp-GTC-3′tDR attenuates cardiac fibrosis and improves cardiac function in the murine MI model

We next investigated the regulation and pathophysiological role of Asp-GTC-3’tDR in the preclinical murine models of cardiac ischemia. Northern blot detected abundant Asp-GTC-3’tDR in the heart tissue 1 and 2 days after MI surgery, with higher levels in the infarct zone (**Figures 2A and S3A**). A similar induction occurred in the murine cardiac I/R model: Asp-GTC-3’tDR levels in ischemic heart tissue rose within 45 minutes of left anterior descending coronary (LAD) artery ligation and remained elevated one day after the I/R surgery (**Figure S3B**). Thus, acute cardiac ischemia enhances Asp-GTC-3’tDR generation in vivo, mirroring its induction in cultured cells. In both models, Asp-GTC-3’tDR levels subsequently declined by one week after surgery (**Figures 2A, S3A-B**), as the acute ischemic phase resolved, indicating that the loss of Asp-GTC-3’tDR biogenesis coincides with the onset of adverse cardiac remodeling.

**Figure 2.**
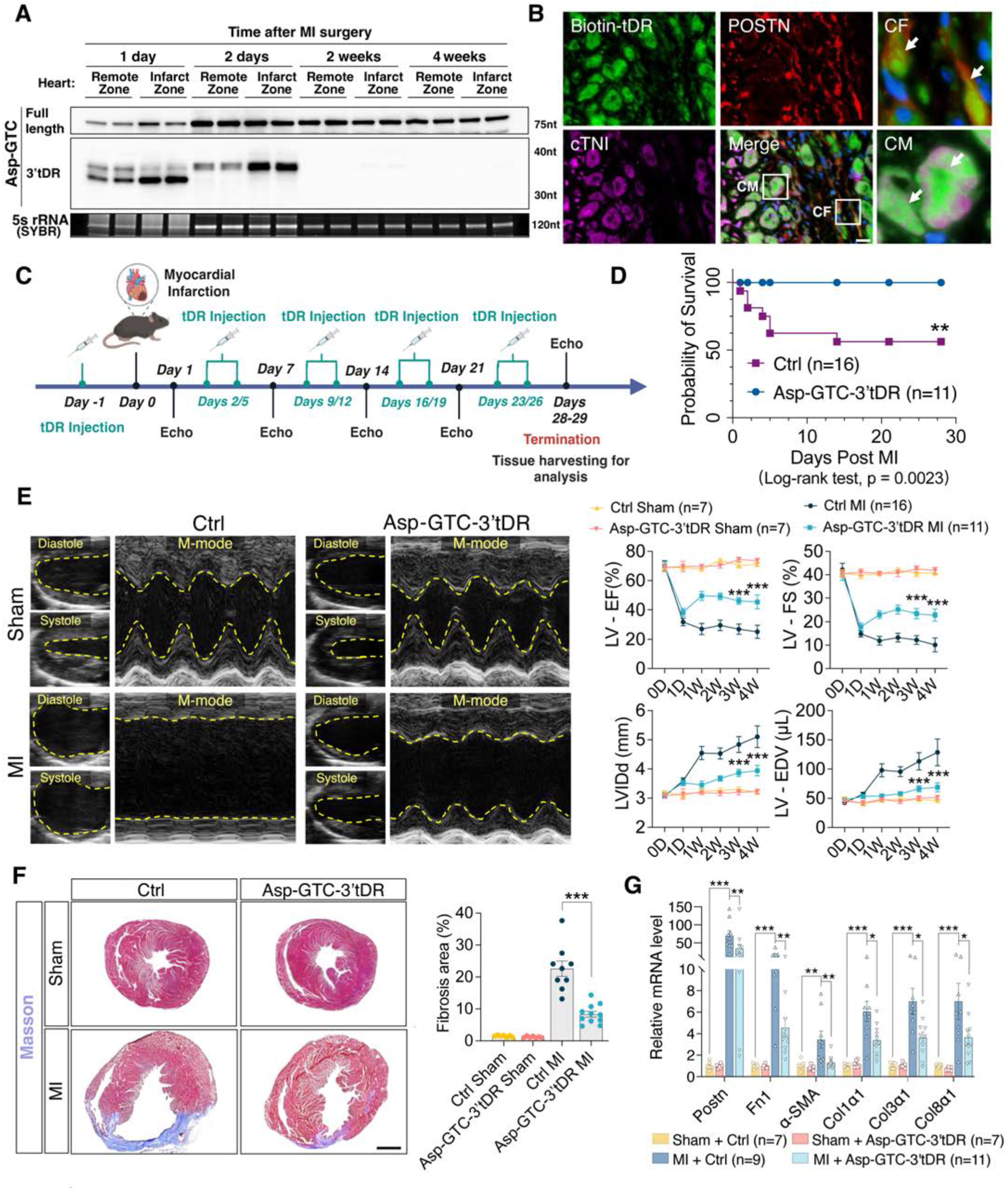
Delivery of Asp-GTC-3′tDR attenuates cardiac remodeling and improves cardiac function in the murine MI model. **(A)** Northern blot analysis of Asp-GTC-3’tDR levels in the remote and infarct zones of mouse hearts following MI surgery. **(B)** Immunofluorescence staining of ischemic heart sections after delivery of PEI-formulated biotin-conjugated Asp-GTC-3’tDR, showing uptake of Asp-GTC-3’tDR (green) in POSTN-positive myofibroblasts (red) and cTNI-positive CMs (magenta). Scale bar, 100 μm. **(C)** Schematic illustration of the experimental design for tDR administration in the murine MI model. **(D)** Kaplan–Meier survival analysis of MI mice treated with Ctrl or Asp-GTC-3′tDR mimics over the 4-week follow-up. **(E)** Representative M-mode echocardiograms (4 weeks after surgery) and quantification of cardiac function, including LV-EF, LV-FS, LV-IDd, and LV-EDV in sham and MI mice treated with Ctrl or Asp-GTC-3’tDR mimics. **(F)** Masson’s trichrome staining and quantification of fibrotic area in heart sections from sham and MI groups treated with Ctrl or Asp-GTC-3’tDR mimics. Scale bar, 2 mm **(G)** qPCR analysis of fibrosis-associated genes in heart tissues from sham and MI mice treated with Ctrl or Asp-GTC-3’tDR mimics. Data are presented as means ± SEM. Statistical significance was determined using the log-rank test (D), two-way ANOVA with appropriate post hoc tests (E), the unpaired two-tailed Student’s t-test (F), or the Mann-Whitney test (G). *\*P* < 0.05, *\*\* P*< 0.01, *\*\*\*P* < 0.001.

To determine if restoring the levels of Asp-GTC-3’tDR could mitigate fibrosis and adverse remodeling after MI, we optimized methods to deliver synthetic tDR mimics to the heart in our preclinical models. We compared two commonly used nonviral vehicles for small RNA delivery and found that the polymer-based polyethyleneimine (PEI) efficiently delivered synthetic tDR mimics to heart tissues with intravenous administration, as confirmed by Northern blot (**Figure S3C**). To test whether PEI could deliver the tDR mimics to CFs in the infarct zone, we administered PEI-formulated biotinylated Asp-GTC-3’tDR mimics to mice 1 week after MI surgery. Fluorescence imaging demonstrated uptake in both CFs and CMs within the infarct zone, indicating effective delivery to key cardiac cell populations (**Figure 2B**). We further assessed the *in vivo* stability of delivered Asp-GTC-3′tDR mimics by monitoring their persistence in heart tissue over time. Northern blot analysis showed that Asp-GTC-3′tDR was readily detectable in the heart at early time points and remained abundant up to 4 days post-injection, indicating that systemically delivered Asp-GTC-3’tDR has sufficient stability and retention in cardiac tissue (**Figure S3D**).

Using our optimized methodology for delivering tDR mimics to the heart, we tested whether augmenting the levels of Asp-GTC-3’tDR is protective after MI. PEI-formulated Asp-GTC-3′tDR or control mimics were administered one day before MI surgery and then twice weekly for 4 weeks (**Figure 2C**). Northern blot analysis confirmed the successful delivery of Asp-GTC-3′tDR mimics to the hearts of treated mice (**Figure S3E**). Notably, Asp-GTC-3′tDR treatment markedly improved post-MI survival compared with control mimics treatment (**Figure 2D**). This survival benefit was associated with a marked improvement in cardiac systolic function. In the control group, cardiac function declined rapidly within 1 day of MI and deteriorated progressively over 28 days without recovery; in contrast, Asp-GTC-3′tDR-treated mice regained left ventricular (LV) systolic function and had amelioration of adverse structural remodeling characterized by significantly higher LV ejection fraction (LV-EF) and fractional shortening (LV-FS), and lower LV internal end-diastolic diameter (LV-IDd) and LV end-diastolic volume (LV-EDV) at 4 weeks (**Figure 2E**). Consistent with improved cardiac remodeling, Asp-GTC-3′tDR-treated mice had a significantly lower heart weight-to-body weight ratio compared with control MI mice (**Figure S3F**). Finally, we assessed cardiac fibrosis by Masson trichrome and Sirius Red staining. MI mice developed severe myocardial fibrosis and collagen deposition in the injured myocardium compared with sham controls, and both were markedly attenuated by Asp-GTC-3′tDR treatment (**Figures 2F and S3G**). In concordance, qPCR confirmed the suppression of fibrosis-associated gene expression in the injured myocardium following Asp-GTC-3′tDR treatment (**Figure 2G**). Collectively, these data demonstrate that Asp-GTC-3′tDR confers robust cardioprotection against post-infarction fibrosis.

### Asp-GTC-3′tDR mitigates pathological cardiac hypertrophy

The beneficial phenotype of Asp-GTC-3’tDR overexpression in the MI model suggests additional protective effects beyond fibrosis. Wheat germ agglutinin (WGA) staining of hearts collected 4 weeks after MI revealed the expected increase in CM cross-sectional area in control-treated mice, which was markedly reduced by Asp-GTC-3′tDR administration (**Figure 3A**). Consistently, transcriptional signatures of pathological hypertrophy, which were significantly induced by MI surgery, were decreased by Asp-GTC-3′tDR treatment (**Figure 3B**). These observations suggested that Asp-GTC-3′tDR may also have a CM-specific effect, restraining pathological hypertrophy, which prompted us to examine its regulation and function in CMs directly. As in CFs, Asp-GTC-3′tDR was strongly induced by ischemia in neonatal rat ventricular CMs (NRVMs) (**Figure 3C**). Conversely, in the phenylephrine (PE)-induced hypertrophy model, Asp-GTC-3′tDR generation was significantly diminished in hypertrophic NRVMs relative to untreated cells (**Figure 3D**). Thus, Asp-GTC-3’tDR shows the same inverse relationship to disease in CMs as in CFs: acute ischemic stress drives a likely compensatory increase, whereas sustained profibrotic or pro-hypertrophic activation depletes it.

**Figure 3.**
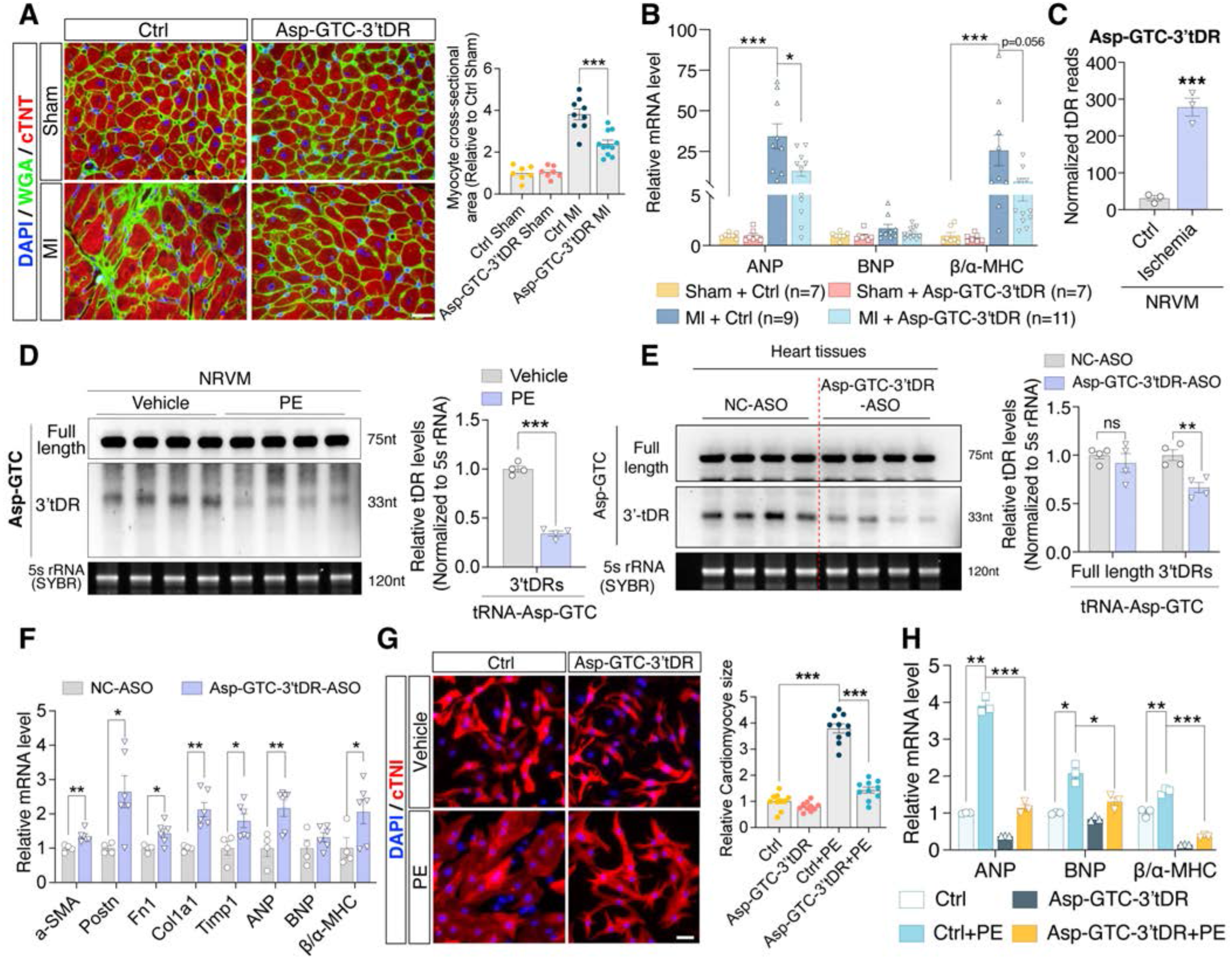
Asp-GTC-3′tDR mitigates pathological cardiac hypertrophy. **(A)** WGA staining and quantification of CM cross-sectional area in heart sections from sham and MI mice treated with Ctrl or Asp-GTC-3′tDR mimics. Scale bar, 20 μm. **(B)** qPCR analysis of hypertrophic markers in cardiac tissues from sham and MI mice treated with Ctrl or Asp-GTC-3′tDR mimics. **(C)** ARM-seq normalized read counts for Asp-GTC-3′tDR in NRVMs with or without 5 hours of ischemia. **(D)** Northern blot analysis and quantification of Asp-GTC-3′tDR levels in NRVMs treated with or without PE for 48 hours. **(E)** Northern blot analysis and quantification of Asp-GTC-3′tDR levels in mouse hearts after treatment with negative control ASO (NC-ASO) or Asp-GTC-3’tDR-ASO for a week. **(F)** qPCR analysis of fibrosis- and hypertrophy-associated genes in mouse hearts after Asp-GTC-3′tDR silencing with ASOs. **(G)** Immunofluorescence staining and quantification of CM cross-sectional area in NRVMs transfected with Ctrl or Asp-GTC-3′tDR mimics and treated with vehicle or PE for 48 hours. Scale bar, 100 μm. **(H)** qPCR analysis of hypertrophic genes in NRVMs transfected with Ctrl or Asp-GTC-3′tDR mimics and treated with vehicle or PE for 48 hours. Data are presented as means ± SEM. Statistical significance was determined using the unpaired two-tailed Student’s t-test (A, C, D, E, and G) or the Mann-Whitney test (B and F). *\*P* < 0.05, *\*\* P*< 0.01, *\*\*\*P* < 0.001.

To test whether endogenous Asp-GTC-3′tDR actively restrains adverse remodeling in the heart, we performed loss-of-function experiments using antisense oligonucleotide (ASO)-mediated silencing. A fully phosphorothioate-modified ASO targeting Asp-GTC-3’tDR (Asp-GTC-3’tDR-ASO^30^) and a negative control ASO (NC-ASO) were administered intravenously into healthy mice every other day for one week. Northern blot confirmed that Asp-GTC-3’tDR-ASO specifically reduced Asp-GTC-3′tDR levels without significantly altering the parent full-length tRNA-Asp-GTC pool (**Figure 3E**). Notably, silencing endogenous Asp-GTC-3′tDR in mouse hearts led to significant increases in both profibrotic and pro-hypertrophic gene expression (**Figure 3F**), indicating that endogenous Asp-GTC-3′tDR tonically counterbalances profibrotic and pro-hypertrophic programs to maintain cardiac homeostasis. To examine its anti-hypertrophic role directly, we transfected NRVMs with control or Asp-GTC-3’tDR mimics before PE stimulation. PE treatment drove a robust increase in CM size in the control group, whereas increased Asp-GTC-3′tDR expression markedly attenuated PE-induced hypertrophy, as evidenced by the reduced CM size and hypertrophic gene expression (**Figures 3G and 3H**), demonstrating a potent anti-hypertrophic function of Asp-GTC-3’tDR. Together with its antifibrotic activity, these results establish Asp-GTC-3′tDR as a multifunctional cardioprotective tDR that coordinately restrains fibrotic and hypertrophic remodeling after cardiac injury.

### Asp-GTC-3′tDR confers therapeutic cardiac benefit in acute and chronic cardiac injury models

The MI experiments established the cardioprotective effects of Asp-GTC-3’tDR when administered prior to ischemic injury. To determine its therapeutic efficacy as a treatment paradigm, we used a cardiac I/R model of acute ischemic injury and a transverse aortic constriction (TAC) model of chronic pressure overload. To model early clinical intervention after an ischemic event, we treated mice with a short treatment duration for the Asp-GTC-3′tDR mimics: a first dose immediately after reperfusion and two booster doses within the first week, followed by an 8-week follow-up (**Figure 4A**). Over this period, I/R injury in control-treated mice produced a rapid and sustained loss of LV systolic function, with decreasing LV-EF and LV-FS, accompanied by progressive chamber dilation (rising LVIDd and LV-EDV). Conversely, Asp-GTC-3’tDR-treated mice demonstrated a significant improvement in LV function and structural remodeling, with improved LV systolic function and chamber dimensions (**Figures 4B and 4C**). The improved function tracked with histological features of reverse remodeling. Control-treated hearts developed extensive interstitial fibrosis and hypertrophy after I/R, whereas Asp-GTC-3′tDR treatment reduced both the fibrotic area and the CM cross-sectional area (**Figures 4D**, **4E, S4A, and S4B**), demonstrating that a limited dosage regimen after reperfusion improved post-ischemic remodeling. These structural benefits were reflected in the transcriptome: Asp-GTC-3′tDR limited the expression of fibrosis-associated genes and hypertrophic markers (**Figure 4F**). Thus, a treatment paradigm of limited Asp-GTC-3′tDR administration following reperfusion attenuated post-ischemic adverse remodeling and drove functional recovery.

**Figure 4.**
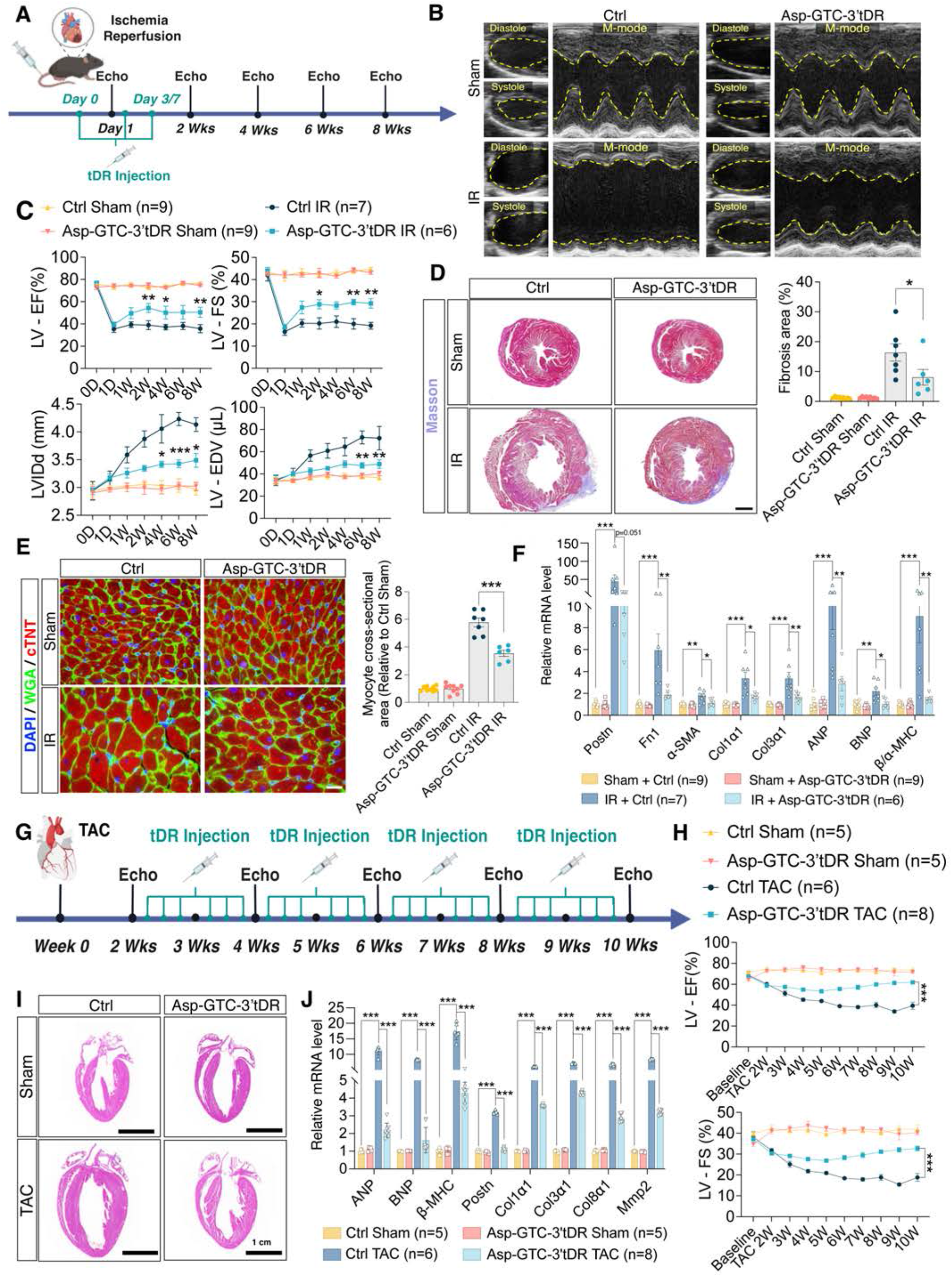
Asp-GTC-3′tDR confers therapeutic cardiac benefit in acute and chronic cardiac injury models. **(A)** Schematic illustration of the experimental design for limited Asp-GTC-3′tDR treatment in the mouse cardiac I/R model. **(B)** Representative M-mode echocardiograms of sham and I/R mice treated with Ctrl or Asp-GTC-3′tDR mimics 8 weeks after surgery. **(C)** Quantification of cardiac function, including LV-EF, LV-FS, LV-IDd, and LV-EDV of sham and I/R mice treated with Ctrl or Asp-GTC-3′tDR mimics. **(D)** Masson’s trichrome staining and quantification of fibrotic area in heart sections from sham and I/R mice treated with control or Asp-GTC-3′tDR mimics. Scale bar, 2 mm. **(E)** WGA staining and quantification of CM cross-sectional area in heart sections from sham and I/R mice treated with Ctrl or Asp-GTC-3′tDR mimics. Scale bar, 20 μm. **(F)** qPCR analysis of fibrosis- and hypertrophy-associated genes in heart tissues from sham and I/R mice treated with Ctrl or Asp-GTC-3’tDR mimics. **(G)** Schematic illustration of the experimental design for Asp-GTC-3′tDR treatment in the mouse TAC model. **(H)** Serial echocardiographic assessment of cardiac function, including LV-EF and LV-FS, in sham or TAC mice treated with Ctrl or Asp-GTC-3′tDR mimics. **(I)** Representative H&E staining of sham and TAC hearts treated with Ctrl or Asp-GTC-3′tDR mimics. Scale bar, 1 cm. **(J)** qPCR analysis of hypertrophy- and fibrosis-associated genes in cardiac tissues from sham and TAC mice treated with Ctrl or Asp-GTC-3′tDR mimics. Data are presented as means ± SEM. Statistical significance was determined using two-way ANOVA with appropriate post hoc tests (C and H), the unpaired two-tailed Student’s t-test (D and E), or the Mann-Whitney test (F and J). *\*P* < 0.05, *\*\* P*< 0.01, *\*\*\*P* < 0.001.

The I/R model tests early intervention, but patients often present after remodeling is established. We therefore turned to the pressure overload model, in which the pathology develops progressively and offers a window for delayed treatment. In the TAC model, we withheld Asp-GTC-3′tDR treatment until two weeks after constriction, by which point hypertrophy and functional decline were already underway, and then dosed the tDR mimics three times weekly (**Figure 4G**), a paradigm that more faithfully models therapeutic intervention in chronic HF. Under these conditions, Asp-GTC-3′tDR administration successfully reversed the trajectory of disease. Cardiac systolic function, which had been declining at the start of treatment, recovered during long-term follow-up, with higher LV-EF and LV-FS than in control-treated mice (**Figure 4H**). This functional rescue was accompanied by beneficial structural remodeling: Asp-GTC-3’tDR-treated hearts were visibly smaller on both gross examination and morphological imaging (**Figures S5A and 4I**), with correspondingly lower heart weight–to–body weight ratios (**Figure S5B**). Interstitial fibrosis was reduced on Masson trichrome staining, and hypertrophic and fibrotic gene expression fell accordingly, in Asp-GTC-3’tDR-treated mouse hearts compared to controls (**Figures S5C and 4J**). Together, these models demonstrated that Asp-GTC-3′tDR is cardioprotective across acute and chronic injury and, critically, across therapeutic windows, effective even when given after the onset of ischemic injury or the establishment of pathological remodeling.

### Asp-GTC-3′tDR attenuates cardiac fibrosis by promoting SG assembly

Our previous work showed that the guanine-rich regions of Asp-GTC-3’tDR facilitated oligo-guanine motif-dependent G-quadruplex (G4) formation, which was essential for its function in kidney cells^30^. We sought to determine whether its antifibrotic function in CFs is similarly structure-dependent. We compared a panel of Asp-GTC-3′tDR mutants, including 12GU, which converts the second guanine in each oligo-guanine motif to uridine; 4CA, which replaces the 4 cytosines in the TΨC stem with adenines; and 12UA, which substitutes the four uridines in the TΨC loop with adenines (**Figure S6A**). Functional screening analysis in CFs revealed that wild-type Asp-GTC-3′tDR suppressed profibrotic gene expression, and 4CA and 12UA mutants retained this activity, whereas only the 12GU mutant was no longer functional (**Figure S6B**). This functional loss tracked with a structural defect: the 12GU mutant failed to form intermolecular G4 structures, whereas the wild-type Asp-GTC-3’tDR and the 4CA and 12UA mutants assembled G4s, forming N-methylmesoporphyrin-positive intermolecular assemblies under G4-permissive (K^+^) but not G4-nonpermissive (Li^+^) conditions (**Figures S6C and S6D**). The failure of the G4-null mutant demonstrates that the G4 structure is specifically required for the antifibrotic function of Asp-GTC-3′tDR.

The function of tDRs often depends on their interactions with proteins^27,28,38,39^, and our prior study demonstrated that Asp-GTC-3’tDR’s function in kidney cells required its binding to PUS7. We therefore characterized the protein partners of Asp-GTC-3′tDR. Biotin-conjugated Asp-GTC-3’tDR or control mimics were transfected into NRCFs, and bound proteins were enriched by streptavidin pulldown and analyzed by proteomics. Interestingly, the interactome was dominated by ribosomal proteins from both small and large subunits, suggesting a role for Asp-GTC-3′tDR in regulating ribosome-related processes (**Figure 5A, Table S2**). Pulldown-immunoblot analysis of the top-ranked candidates also confirmed the strong interaction between Asp-GTC-3’tDR and ribosomal proteins (**Figure 5B**). This association with the translational machinery initially suggested that Asp-GTC-3′tDR may repress global translation by competing with mRNAs for ribosome binding, as reported for other tDRs^40^. However, the puromycin-incorporation assay showed no change in bulk protein synthesis between control- and Asp-GTC-3′tDR-treated NRCFs (**Figure 5C**). Therefore, Asp-GTC-3′tDR associates with ribosomes without acting as a global translational repressor, suggesting selective regulation of ribosome-associated pathways.

**Figure 5.**
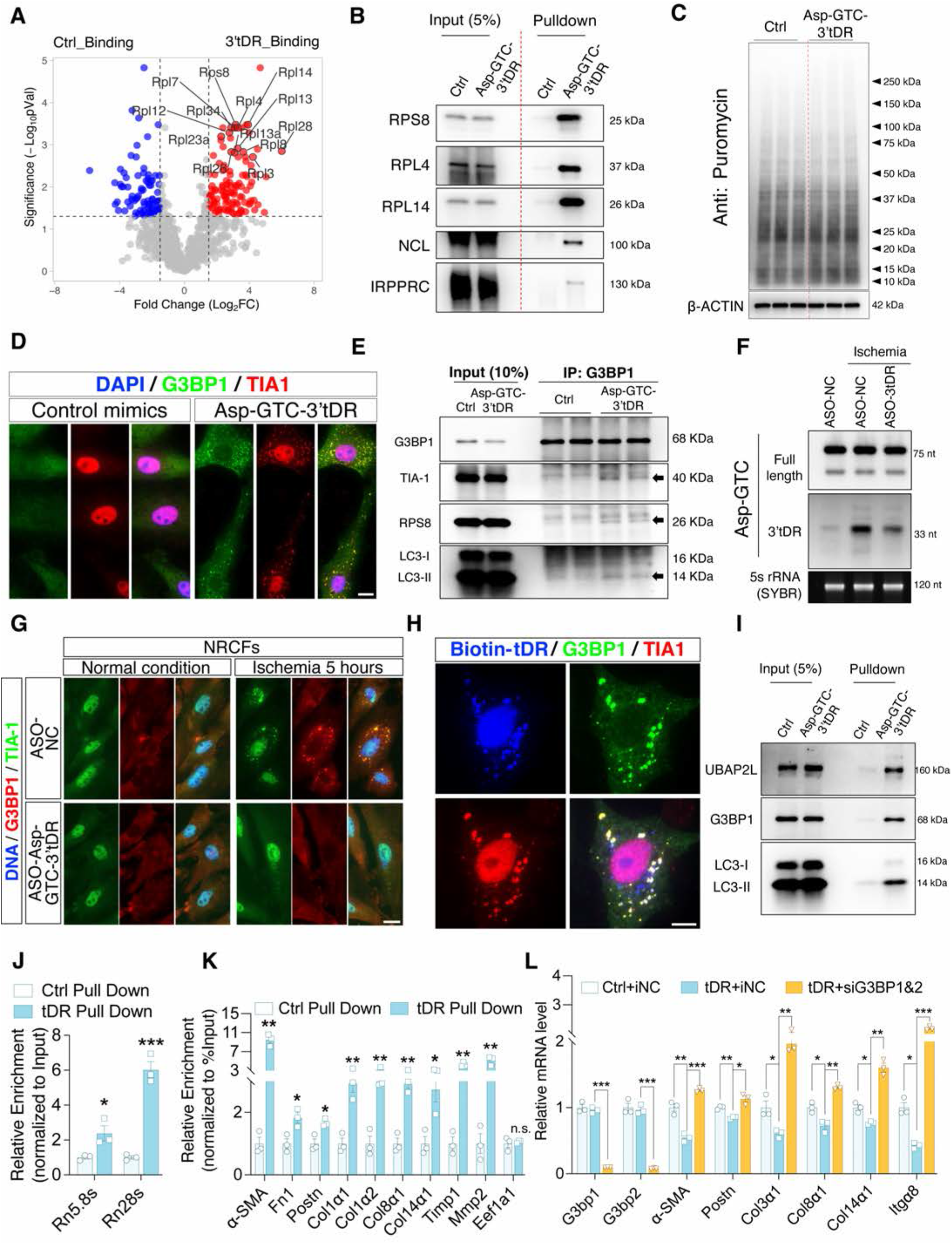
Asp-GTC-3′tDR attenuates cardiac fibrosis by promoting SG assembly. **(A)** Proteomic analysis of proteins associated with Asp-GTC-3′tDR, after pulldown of biotinylated Ctrl or Asp-GTC-3’tDR mimics from NRCFs using streptavidin beads. Significant binding candidates (P < 0.05) are highlighted in red (log2[fold change] >1) or blue (log2[fold change] < –1). **(B)** Immunoblot validation of the top 5 Asp-GTC-3′tDR-associated proteins after streptavidin pulldown of biotinylated Ctrl or Asp-GTC-3’tDR mimics from NRCFs. **(C)** Puromycin incorporation assay of global protein synthesis in NRCFs 24 hours after transfection of Ctrl or Asp-GTC-3′tDR mimics. **(D)** Representative immunofluorescence of G3BP1- and TIA1-positive SGs in NRCFs 24 hours after transfection of Ctrl or Asp-GTC-3′tDR mimics. Scale bar, 20 μm. **(E)** Immunoprecipitation of G3BP1 co-isolated the SG component TIA1, the ribosomal protein RPS8, and the autophagy protein LC3 in NRCFs transfected with Asp-GTC-3′tDR mimics but not Ctrl mimics. **(F)** Northern blot analysis of endogenous Asp-GTC-3′tDR levels following ASO-mediated silencing in NRCFs under ischemia. (**G**) Representative immunofluorescence of SG formation in ischemic NRCFs following Asp-GTC-3′tDR silencing. Scale bar, 20 μm. (**H**) Representative fluorescence images showing colocalization of biotinylated Asp-GTC-3′tDR (blue) with G3BP1- and TIA1-positive SGs in NRCFs. Scale bar, 20 μm. **(I)** Asp-GTC-3’tDR pulldown followed by immunoblot of the SG-associated proteins UBAP2L and G3BP1 and the autophagy protein LC3-II in NRCFs transfected with biotinylated Ctrl or Asp-GTC-3’tDR mimics. **(J)** qPCR analysis of 5.8S and 28S rRNA levels enriched by Asp-GTC-3′tDR pulldown relative to Ctrl pulldown in NRCFs transfected with biotinylated Ctrl or Asp-GTC-3’tDR mimics. **(K)** qPCR analysis of profibrotic transcripts enriched by Asp-GTC-3′tDR pulldown relative to Ctrl pulldown in NRCFs transfected with biotinylated Ctrl or Asp-GTC-3’tDR mimics. Enrichment was normalized to input and expressed relative to the Ctrl pulldown. **(L)** qPCR analysis of profibrotic gene expression in NRCFs transfected with Ctrl or Asp-GTC-3′tDR mimic after co-silencing of G3BP1 and G3BP2 (siG3BP1&2). Data are presented as means ± SEM. Statistical significance was determined using the unpaired two-tailed Student’s t-test. n.s., nonsignificant. *\*P* < 0.05, *\*\* P*< 0.01, *\*\*\*P* < 0.001.

Selective translational control that spares bulk protein synthesis is a defining property of SGs, cytoplasmic condensates that nucleate from stalled initiation complexes and transiently sequester non-translating mRNAs during stress ^26^. Consistent with a role in this process, Asp-GTC-3′tDR overexpression robustly induced the formation of G3BP1- and TIA1-positive cytoplasmic granules in cultured neonatal and adult CFs and in the fibrosis regions of MI hearts (**Figures 5D, S7A, and S7B**). Moreover, immunoprecipitation of G3BP1 showed increased association of core SG components and ribosomal proteins in Asp-GTC-3′tDR-treated CFs compared to controls (**Figure 5E**), together supporting that Asp-GTC-3’tDR enhances SG assembly in CFs. The same immunoprecipitation also demonstrated a higher association of lipidated LC3 proteins with G3BP1 after Asp-GTC-3′tDR treatment (**Figure 5E**). Because unresolved SGs are typically cleared through autophagy ^41^, this recruitment of autophagy machinery suggests that Asp-GTC-3′tDR-nucleated SGs are likely routed for autophagic degradation. Next, we asked whether endogenous Asp-GTC-3′tDR drives SG formation. Acute ischemia treatment enhanced SG assembly in NC-ASO-transfected NRCFs, whereas ASO-mediated silencing of Asp-GTC-3′tDR reduced G3BP1/TIA1-positive SGs in NRCFs under the ischemic condition (**Figures 5F and 5G**). Thus, these findings demonstrate that Asp-GTC-3′tDR was both sufficient and necessary for promotion of SG assembly in CFs during ischemic stress.

To further probe the role of Asp-GTC-3’tDR in SG assembly, we tracked the subcellular localization of biotinylated Asp-GTC-3′tDR and observed extensive colocalization with G3BP1- and TIA1-positive SGs in NRCFs (**Figure 5H**). Consistently, pulldown of the biotinylated Asp-GTC-3’tDR recovered core SG components, including G3BP1 and UBAP2L, together with the autophagic protein LC3-II (**Figure 5I**). The same pulldown also enriched 5.8S and 28S rRNAs (**Figure 5J**), consistent with the ribosomal-protein binding above and indicating that Asp-GTC-3′tDR engages SGs and ribosomes simultaneously. These results place Asp-GTC-3′tDR physically within SGs.

Because Asp-GTC-3’tDR lowers mRNA levels of profibrotic genes, we asked whether these profibrotic transcripts are selectively recruited into SGs for translational silencing and subsequent autophagic clearance. Pulldown of biotinylated Asp-GTC-3’tDR enriched multiple profibrotic transcripts but not the housekeeping Eef1a1 mRNA (**Figure 5K**), indicating that these ribosome-loaded profibrotic transcripts are selectively recruited into Asp-GTC-3’tDR-associated SGs. Together with the reduced fibrotic mRNA levels, decreased collagen synthesis, and the recruitment of autophagy machinery, these observations suggest that Asp-GTC-3’tDR partitions profibrotic transcripts into SGs for translation silencing and subsequent clearance. To test whether SG formation is required rather than merely correlated, we blocked SG assembly by simultaneously silencing the nucleators G3BP1 and G3BP2 ^42,43^. In the absence of G3BP1 and G3BP2, Asp-GTC-3’tDR could no longer promote SG formation or suppress profibrotic gene expression (**Figures S7C and 5L**), establishing that SG assembly is mechanistically required for its antifibrotic activity. Together, these findings define a mode of tDR-mediated gene regulation in CFs: rather than repressing global translation, Asp-GTC-3′tDR nucleates SGs and facilitates the partitioning of profibrotic transcripts into SGs, restraining fibrotic gene expression and thereby ameliorating pathological myocardial fibrosis.

### Asp-GTC-3′tDR sequesters PUS7 to enhance RNA autophagy in CMs

We next asked whether the same mechanism of SG nucleation and mRNA portioning observed in fibroblasts underlies its antihypertrophic effects in CMs. Interestingly, immunofluorescence analysis revealed that Asp-GTC-3′tDR overexpression failed to induce G3BP1- and TIA1-positive SG formation in NRVMs (**Figure 6A**), and no increase in G3BP1-positive granules was observed in CMs in heart tissues from Asp-GTC-3′tDR-treated mice (**Figure S8A**). These results suggest that Asp-GTC-3′tDR acts through a cell-type-specific mechanism, engaging a process in CMs distinct from the SG-based pathway in CFs. To identify that pathway, we profiled the transcriptomic changes driven by Asp-GTC-3’tDR in CMs. Hypertrophic NRVMs were transfected with control or Asp-GTC-3’tDR mimics and analyzed by RNA sequencing. The transcriptome analysis revealed that, beyond the expected suppression of hypertrophic genes, Asp-GTC-3’tDR downregulated a specific set of histone mRNAs (**Figure 6B**), which qPCR confirmed across multiple histone transcripts (**Figure 6C**), thereby identifying histone mRNAs as a distinct class of Asp-GTC-3’tDR targets in CMs. This was mechanistically similar to our findings in kidney cells, where Asp-GTC-3’tDR sequesters PUS7 from its histone mRNA substrates, and the resulting pseudouridylation-deficient histone mRNAs are routed to the autophagosome/lysosome pathway for degradation, a process termed RNA autophagy ^30^. Consistent with the same axis operating in CMs, pulldown of biotinylated Asp-GTC-3’tDR from NRVMs recovered PUS7, along with PUS1 and DHX36 (**Figure 6D**), the same interactors we identified in kidney cells^30^. We therefore tested whether Asp-GTC-3’tDR engages this PUS7-RNA autophagy axis to control histone mRNAs and thereby hypertrophy, in CM.

**Figure 6.**
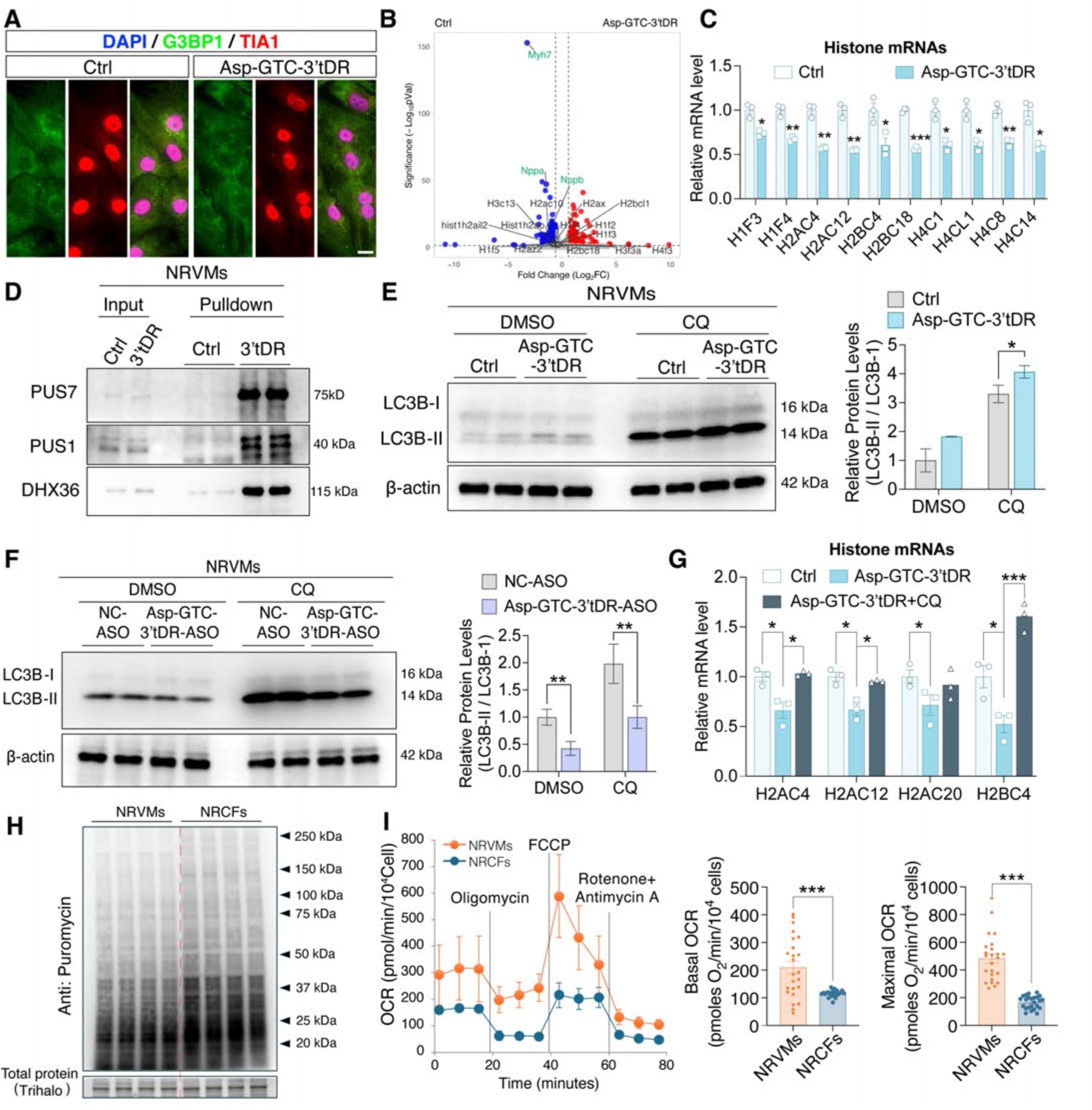
Asp-GTC-3′tDR sequesters PUS7 to enhance RNA autophagy in CMs. **(A)** Representative immunofluorescence of G3BP1- and TIA1-positive SGs in NRVMs 24 hours after transfection of Ctrl or Asp-GTC-3′tDR mimics. Scale bar, 20 μm. **(B)** Volcano plot of differentially expressed genes in hypertrophic NRVMs transfected with Asp-GTC-3′tDR mimics versus Ctrl mimic. **(C)** qPCR analysis of histone mRNAs in NRVMs transfected with Ctrl and Asp-GTC-3′tDR mimics. **(D)** Asp-GTC-3’tDR pulldown followed by immunoblot analysis of PUS7, PUS1, and DHX36 in NRVMs transfected with biotinylated Ctrl or Asp-GTC-3’tDR mimics. **(E)** Immunoblot analysis and quantification of the LC3B-II/LC3B-I ratio in NRVMs transfected with Ctrl or Asp-GTC-3′tDR mimic and treated with dimethyl sulfoxide (DMSO) or CQ for 2 hours before collection. **(F)** Immunoblot analysis and quantification of LC3B-II/LC3B-I ratio in NRVMs transfected with NC-ASO or Asp-GTC-3′tDR-targeting ASO and treated with DMSO or CQ for 2 hours before collection. **(G)** qPCR analysis of histone mRNAs in NRVMs transfected with Ctrl or Asp-GTC-3′tDR mimics and treated with or without CQ for 2 hours before collection. **(H)** Puromycin incorporation assay comparing global protein synthesis between NRVMs and NRCFs. Total protein **(I)** Seahorse extracellular flux analysis comparing mitochondrial respiration in NRVMs and NRCFs. Oxygen consumption rate (OCR) was measured at baseline and after sequential addition of oligomycin, FCCP, and rotenone plus antimycin A; basal and maximal OCR are quantified. Data are presented as means ± SEM. Statistical significance was determined using the unpaired two-tailed Student’s t-test. *\*P* < 0.05, *\*\* P*< 0.01, *\*\*\*P* < 0.001.

Immunoblot analysis showed that Asp-GTC-3′tDR significantly increased lipidated LC3B-II levels in NRVMs in the presence of chloroquine (CQ) (**Figure 6E**), which blocks lysosomal acidification and enables measurement of autophagic flux^44^. Conversely, ASO-mediated silencing of endogenous Asp-GTC-3′tDR lowered lipidated LC3B-II levels under both basal and CQ-treated conditions (**Figure 6F**), providing confirmation that Asp-GTC-3′tDR promotes autophagic flux in CMs. We then asked whether Asp-GTC-3’tDR activates histone mRNA autophagy in CMs, in parallel to our observations in kidney cells. NRVMs were transfected with Asp-GTC-3’tDR and then treated with CQ for 2 hours to block lysosomal degradation prior to RNA isolation. Expectedly, Asp-GTC-3’tDR leads to a decrease in the levels of key histone mRNAs, while blocking lysosomal degradation with CQ largely abolished this reduction (**Figure 6G**), demonstrating that autophagic flux driven by Asp-GTC-3′tDR in CMs involves histone mRNA autophagy. Notably, the species of histone mRNAs affected by Asp-GTC-3’tDR in rat CMs were largely those we described in kidney cells and carried multiple PUS7-preferred UNUAR consensus motifs (**Table S3**), marking them as substrates for PUS7-dependent pseudouridylation. Together, these results show that Asp-GTC-3′tDR does not form SGs in CMs but instead triggers the PUS7-dependent RNA autophagy pathway, mirroring the mechanism we established in the kidney and defining a second, cell-type-specific route by which this tDR restrains pathological remodeling.

To understand what directs the same tDR toward different pathways in the two cell types, we compared the machineries for SG assembly and RNA autophagy, together with the cells’ physiological demands, between CFs and CMs. The two effector pathways were reciprocally poised. CFs expressed higher levels of cytoplasmic ribosomal proteins, including RPS8 and RPL14, whereas CMs expressed comparatively less ribosomal protein but more PUS7, along with higher glycolytic GAPDH (**Figure S8B**). These abundances tracked with the cells’ functional states: puromycin-incorporation assays showed a higher global translational rate in cultured CFs than in CMs (**Figure 6H**), and Seahorse analysis revealed higher mitochondrial oxidative phosphorylation activity in CMs than in CFs (**Figures 6I**). Thus, each cell type provides an intracellular environment matched to a different arm of Asp-GTC-3′tDR function: abundant ribosomes and a high translational rate favor SG-based translational regulation in fibroblasts, whereas elevated PUS7 and high metabolic demand favor PUS7-dependent RNA autophagy in CMs. This cellular context, rather than a difference in the tDR itself, appears to determine which protective program is deployed by the tDR in response to similar stress signals.

### The cell-type-specific functions of Asp-GTC-3′tDR are conserved in human cardiac cells and disease

Having defined two distinct protective mechanisms in rodent CFs and CMs, we asked whether both are conserved in human cardiac cells. In primary human adult CFs, Asp-GTC-3′tDR reproduced its rodent antifibrotic activity: it significantly suppressed profibrotic gene expression under basal conditions and attenuated the TGFβ-induced activation of fibrosis-associated genes (**Figures 7A and 7B**), and it induced G3BP1- and TIA1-positive SGs (**Figure 7C**), indicating that the same SG-based mechanism operates in human CFs. Similarly, in human iPSC-derived cardiac organoids, Asp-GTC-3′tDR suppressed the expression of endothelin-1 (ET-1)-induced hypertrophic markers (**Figure 7D**), and in human iPSC-derived CMs, it reduced multiple histone transcripts (**Figure 7E**), recapitulating the PUS7/histone mRNA axis defined in rodents. Thus, both cell-type-specific programs, SG-based control in CFs and histone mRNA regulation in CMs, are preserved in human cardiac cells.

**Figure 7.**
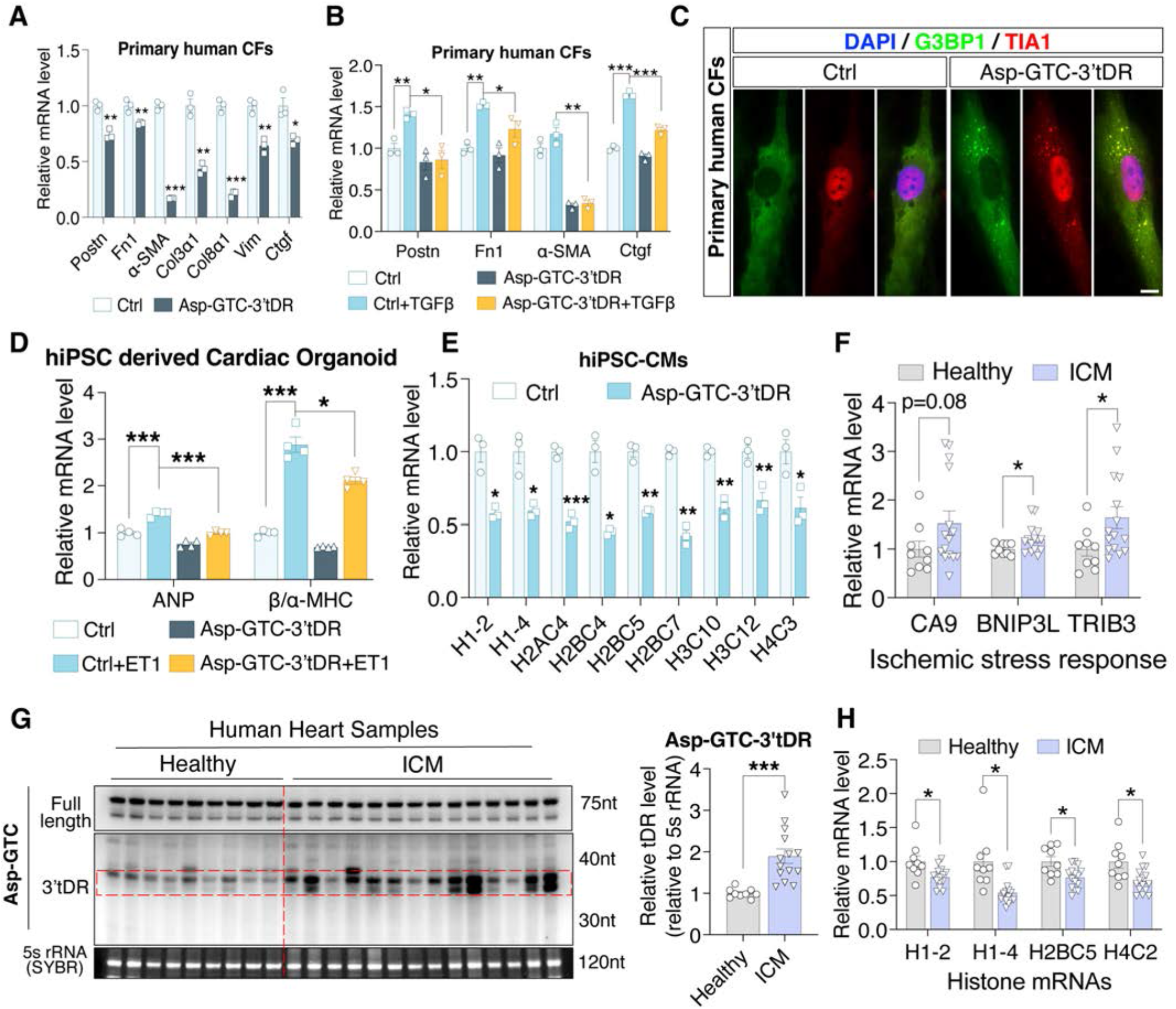
The cell-type-specific functions of Asp-GTC-3′tDR are conserved in human cardiac cells and disease. **(A)** qPCR analysis of profibrotic genes in primary adult human CFs 24 hours after transfection with Ctrl or Asp-GTC-3′tDR mimics. **(B)** qPCR analysis of fibrosis-associated genes in primary adult human CFs transfected with Ctrl or Asp-GTC-3′tDR mimic and treated with vehicle or TGF-β for 48 hours. **(C)** Representative immunofluorescence of G3BP1- and TIA1-positive SGs in primary adult human CFs 24 hours after transfection with Ctrl or Asp-GTC-3′tDR mimic. Scale bar, 20 μm. **(D)** qPCR analysis of hypertrophic genes in human iPSC-derived cardiac organoids transfected with Ctrl or Asp-GTC-3′tDR mimic and treated with or without ET-1 for 48 hours. **(E)** qPCR analysis of histone mRNAs in human iPSC-derived CMs (hiPSC-CMs) treated with Ctrl or Asp-GTC-3′tDR mimic. **(F)** qPCR analysis of chronic ischemic stress-response genes in human heart tissues from healthy donors and patients with ICM. **(G)** Northern blot analysis and quantification of Asp-GTC-3′tDR levels in human heart tissues from healthy donors and patients with ICM. **(H)** qPCR analysis of histone mRNAs in human heart tissues from healthy donors and patients with ICM. Data are presented as mean ± SEM. Statistical significance was determined using the unpaired two-tailed Student’s t-test (A, B, D, and E) or the Mann-Whitney test (F, G, and H). *\*P* < 0.05, *\*\* P*< 0.01, *\*\*\*P* < 0.001.

We next asked whether this protective axis is engaged in human heart disease, using cardiac tissue from healthy donors and patients with ischemic cardiomyopathy (ICM). ICM hearts showed hallmarks of chronic ischemic stress, with elevated CA9, BNIP3L, and TRIB3 (**Figure 7F**), the physiological setting expected to induce Asp-GTC-3′tDR. Consistent with this, Northern blot revealed significantly higher Asp-GTC-3′tDR levels in ICM hearts than in healthy controls (**Figure 7G**). Critically, the downstream signature of the tDR was also present: histone mRNAs were reduced in ICM hearts relative to healthy controls (**Figure 7H**), mirroring in patients the tDR/histone mRNA autophagy relationship we defined mechanistically in CMs. These data validate the human relevance of our findings, demonstrating the induction of Asp-GTC-3’tDR by chronic ischemic stress in human ventricular tissues. This is accompanied by the suppression of its histone mRNA targets, providing evidence that the endogenous protective response we characterized is activated in human disease. Together, these results establish that both functional arms of Asp-GTC-3′tDR are conserved in human cardiac cells and that the tDR/histone mRNA autophagy axis operates in human ischemic heart disease, supporting the translational potential of Asp-GTC-3′tDR as a regulator of pathological cardiac remodeling.

## DISCUSSION

Emerging evidence has recognized tDRs as evolutionarily conserved small non-coding RNA regulators of cellular adaptation to stress, orchestrating diverse biological processes that range from global translation to miRNA-like silencing and autophagy^19,21^. Here, we identify Asp-GTC-3′tDR as a stress-induced tDR that protects the heart against pathological remodeling through distinct cell-type-specific mechanisms. Asp-GTC-3′tDR is generated from tRNA-Asp-GTC by ANG-mediated cleavage acutely after ischemic injury. While initially adaptive and cardioprotective, ‘exhaustion’ of tDR biogenesis during the later chronic phase leads to declining expression levels that temporally coincide with myofibroblast activation and CM hypertrophy. Restoring Asp-GTC-3’tDR suppresses both pathological fibrosis and hypertrophy in multiple preclinical HF models, whereas depleting it is sufficient to induce these adverse remodeling programs in healthy hearts. Mechanistically, the same tDR nucleates SGs to restrain profibrotic gene expression in CFs but sequesters PUS7 to drive RNA autophagy and attenuate hypertrophy in CMs.

The importance of these findings should be interpreted in the context of the substantial residual risk of poor clinical outcomes and progressive adverse structural ventricular remodeling in patients with HF, including those receiving GDMT ^6,7^. This observation suggests that key maladaptive remodeling pathways, particularly cardiac fibrosis and hypertrophy, are not directly addressed by current therapies. Therefore, identifying the molecular regulators that govern these processes may reveal therapeutic targets complementary to GDMT. Our work establishes Asp-GTC-3′tDR as a key regulator of cardiac remodeling and targeting it with a synthetic mimic ameliorated both fibrosis and hypertrophy across multiple murine HF models, through a mechanism distinct from the neurohormonal pathways targeted by current therapy. More broadly, these findings extend the biology of tDRs by demonstrating the importance of cellular context for the mechanistic basis of tDR function.

tDRs have been abundantly detected in hypertrophic and failing myocardium and in the circulation ^45^, where they have shown promise as biomarkers. Although a few studies have reported the miRNA-like function of tDRs in cultured CM models ^46,47^, their pathophysiological roles in cardiac cells or in animal models of cardiovascular diseases remain largely unexplored. Against this backdrop, our study advances the field on three fronts. Firstly, Asp-GTC-3′tDR is not merely a bystander induced by ischemic injury but appears to play an important homeostatic role in the heart, serving as a counter-regulator against pro-fibrotic and pro-hypertrophic pathways. Accordingly, its depletion in the uninjured heart, using the ASO designed to specifically silence the tDR, provoked fibrotic and hypertrophic programs. Secondly, restoring or enhancing Asp-GTC-3’tDR levels by delivering synthetic tDR mimics was protective across a variety of mechanistically distinct HF models spanning both ischemic and non-ischemic remodeling, providing proof of concept for drugging a previously intractable molecular target in the heart. Thirdly, the mechanisms of Asp-GTC-3’tDR’s cardioprotection are grounded in distinct cell-type-specific molecular interactions, including association with ribosomes and nucleation of ribosome-loaded profibrotic mRNAs into SGs in CFs, and sequestration of PUS7 to activate RNA autophagy (and in turn bulk autophagy) in CMs. Together, these results suggest that the broader repertoire of stress-induced cardiac tDRs may harbor additional regulators of remodeling, an area that remains largely uncharted.

The therapeutic efficacy of Asp-GTC-3′tDR is notable across our three different preclinical HF models. Augmentation of Asp-GTC-3’tDR levels by delivering synthetic mimics attenuated pathological fibrosis and hypertrophy, improved cardiac function, and improved survival after MI. Delivery of the tDR mimics also conferred similar benefits in cardiac I/R and pressure-overload models, covering both ischemic and non-ischemic etiologies. Two features are particularly relevant for translation. First, in the pressure-overload model, treatment initiated two weeks after banding, once pathological hypertrophy was established, produced a robust improvement in cardiac function, indicating efficacy as a therapeutic intervention rather than only as prophylaxis. Second, in the I/R model, three doses delivered in the first week after reperfusion produced durable benefit over an eight-week follow-up, suggesting that limited augmentation of Asp-GTC-3’tDR expression early after reperfusion can have lasting beneficial effects. Importantly, early suppression of fibrosis by Asp-GTC-3’tDR after ischemic injury carried no adverse consequences, with no evidence of cardiac rupture. This reflects attenuation, rather than complete ablation, of the fibrotic response, consistent with previous studies showing that inhibiting myofibroblast activation pathways during the acute phase of cardiac ischemic injury confers benefit without adverse consequences ^48,49^. We note that the delivery vehicle used here, PEI, is immunogenic, and our mimics lack the modifications needed for enhanced metabolic stability; as such, they are not ready for clinical use. However, as a short structured RNA, Asp-GTC-3’tDR is amenable to chemical engineering, and established stabilization chemistries and targeting conjugates could be combined to achieve in vivo stability and cardiac-selective delivery while preserving its structural motifs and cardioprotective functions.

Our results identified the mechanistic underpinning of tDR engagement with the SG pathway. Prior studies in cultured U2OS cells demonstrated that the 5′ halves of tRNA-Ala-AGC and tRNA-Cys-GCA, which carry 5’ terminal oligo-guanine motifs, assemble into G4 structures and nucleate SGs ^28,50^, an activity linked to neuronal protection ^29^. However, SG formation by these 5’ tRNA halves was demonstrated in a single osteosarcoma cell line, U2OS, and whether SG formation is required for the associated neuronal protection was not defined. We found that Asp-GTC-3′tDR, a 3′ tDR bearing internal rather than terminal oligo-guanine motifs, likewise formed G4 structure and nucleated SGs in CFs. We further showed that the resulting SGs selectively captured ribosome-loaded profibrotic transcripts, thereby restraining their translation and lowering their mRNA abundance, likely due to the autophagic clearance of SGs. This transcript-selective sequestration is genetically required for the antifibrotic effect, as co-depletion of G3BP1 and G3BP2 abolished both SG formation and the antifibrotic effect of Asp-GTC-3’tDR. Together, these findings provide a mechanistic account of how tDR-induced SGs confer protection and identify a protective function for SGs in the heart, consistent with the antifibrotic effects of G3BP1 overexpression^51^. While we did not conclusively determine why transcripts associated with myofibroblast activation are selectively partitioned into the SGs, we speculate that this simply reflects stoichiometry: myofibroblast activation is accompanied by a marked increase in the translation of profibrotic genes ^52^, rendering them the predominant ribosome-loaded species available for capture. Notably, this SG-based mechanism is cell-type-specific: Asp-GTC-3′tDR did not appreciably induce SGs in CMs, indicating that the same tDR does not default to granule formation wherever it is expressed.

In CMs, Asp-GTC-3′tDR did not induce SGs but instead sequestered PUS7 from its histone mRNA substrates, which mechanistically mirrors our findings in kidney cells ^30^; the resulting pseudouridylation-deficient transcripts were routed to the autophagosome/lysosome for degradation, thereby activating RNA autophagy. Importantly, according to our previous work^30^, this selective histone mRNA autophagy did not alter histone protein homeostasis, indicating that the autophagic processing of these pseudouridylation-deficient mRNAs serves not to deplete histones but to act as a trigger that enhances bulk autophagic flux. The recurrence of this axis in a second organ fits its tropism for metabolically active tissues: like the kidney, the heart sustains high basal autophagy to meet its metabolic demand, and Asp-GTC-3′tDR is abundant at baseline in precisely these tissues, positioning it to reinforce autophagic capacity where the pathway is most heavily relied upon. Autophagy activation is itself an established antihypertrophic mechanism ^53^, and the increased autophagic flux we observe is consistent with the reduced hypertrophy conferred by the tDR. This mechanism distinguishes the CM arm from the SG-based route in CFs and suggests that the PUS7/RNA autophagy regulatory function of Asp-GTC-3′tDR may be deployed where high metabolic demand places a premium on autophagic capacity. Consistent with this, our results demonstrated that CMs exhibited higher baseline metabolic activity, higher cytoplasmic PUS7 protein levels, and lower cytoplasmic ribosomal protein levels than CFs. In contrast, CFs exhibit lower baseline metabolic activity, and the ratio of ribosomal proteins to PUS7 is reversed in the cytoplasm. The abundance of cytoplasmic PUS7 in CMs is notable, given that PUS7’s access to cytoplasmic mRNA substrates is a key determinant of its activity ^54^; a larger cytoplasmic PUS7 pool provides more enzyme for Asp-GTC-3’tDR to sequester away from its histone mRNA targets.

That the same tDR executes two distinct programs depending on the cellular context has not been previously reported for this class of RNA, but it echoes a theme emerging for other regulatory RNAs. Individual microRNAs can exert opposite effects in different cardiac cells. miR-30d, for instance, is protective in CMs yet signals to fibroblasts through distinct routes ^55^, and context-dependent outputs have been described for long noncoding RNAs and circular RNAs ^56,57^. Our data extend this principle to tDRs and offer a concrete molecular basis for it. Asp-GTC-3′tDR is routed to SG assembly or to PUS7-linked autophagy according to the effectors each cell type makes available in proximity to the tDR. The high translational load during CF activation makes them particularly susceptible to SG formation, whereas the high metabolic demand in CMs and kidney cells makes these cells more sensitive to the regulation of autophagy. We propose that the stoichiometry (i.e., relative abundance of these effectors, PUS7 or ribosomes) in the relevant subcellular compartment sets that program the tDR executes and could serve as a lever to bias tDR function. While we consider this to be the most likely mechanism for these cell-specific tDR functions, we were not able to rigorously test this experimentally, due to the lack of appropriate reagents and tools.

Finally, our human data indicate the relevance of tDR regulation and function in human HF. In ICM hearts, which bore the molecular signatures of ongoing ischemic stress, Asp-GTC-3′tDR was elevated relative to healthy myocardium, and its histone mRNA targets were correspondingly reduced, mirroring in patient tissue the tDR/RNA autophagy relationship we defined mechanistically in our preclinical murine and cellular models. Its antifibrotic and antihypertrophic actions were also reproduced in primary human adult CFs and human iPSC-derived cardiac organoids. Together, these observations suggest that the endogenous protective response we characterize in rodents is likely operative in human ischemic heart disease and support the relevance of Asp-GTC-3′tDR as a therapeutic target.

### Limitations of the study

There are several limitations in this study. First, the synthetic tDR mimics used in our experiments lack the base modifications and terminal structures of endogenously generated tDRs, due to technical constraints and the poorly characterized modification patterns of these tDRs. Reassuringly, our data indicate that the key function of Asp-GTC-3’tDR depends on its G4 structures rather than on base modifications, and our ASO-based loss-of-function experiments, which act on the endogenous tDRs, support the conclusion drawn from the mimics. Second, although Asp-GTC-3′tDR enhances autophagy and restrains hypertrophy in CMs, we have not resolved whether this protection derives specifically from PUS7-dependent RNA autophagy or from the bulk autophagic flux it triggers. Because RNA autophagy and general autophagy share the same downstream machinery and are closely coupled, selectively attributing the antihypertrophic effect to one or the other is experimentally difficult and remains to be addressed. Third, our proposal that effector abundance directs the same tDR toward SGs or autophagy is based on correlative differences between CFs and CMs; the decisive test, altering the effector balance to redirect the output, remains to be performed, as does direct proof that SG-sequestered transcripts are cleared by autophagy. Relatedly, although existing literature supports a prominent role for autophagy in metabolically active cells and for SGs in cells with bursts of high translational activity ^58,59^, we did not experimentally test this hypothesis. Finally, while the therapeutic delivery of Asp-GTC-3’-tDR was beneficial across several pre-clinical cardiovascular disease models, in both preventative and therapeutic paradigms, treatments were initiated relatively early in the disease trajectory (at the time of revascularization or two weeks after TAC). Whether this strategy remains efficacious when the disease is more established with overt fibrosis and more significant cardiac dysfunction has not yet been addressed. In addition, whether Asp-GTC-3’tDR treatment would have additive or synergistic effects with current GDMT in these preclinical models may be of importance to establish prior to clinical translation.

## Resource availability

### Lead contact

Further information and requests for resources and reagents should be directed to and will be fulfilled by the lead contact, Dr. Guoping Li.

### Materials availability

This study did not generate new, unique reagents.

### Data and code availability

The ARM-seq data of cardiac cells treated with different stressors were obtained from GSE173806 (https://www.ncbi.nlm.nih.gov/geo/query/acc.cgi?acc=GSE173806). Proteomics data were deposited in PRIDE (accession number: PXD082983). Other materials are available from Dr. Guoping Li under a material transfer agreement with Massachusetts General Brigham.

## Acknowledgements

This work was supported by the National Institutes of Health (NIH) (R01HL174709) and the American Heart Association (AHA) Career Development Award (23CDA1045944) awarded to G.L. and the NIH (R35 HL150807) award to S.D. G.L. was also supported by NIH (R01 HL188041), and he is a member of the Human RNome Consortium. T.H. was supported by the AHA Postdoctoral Fellowship (26POST1553125). P.G. was supported by the AHA Postdoctoral Fellowship (23POST1014230). E.C. is supported by MGH-ECOR-FMD (GR1000237) and the AHA career development award (25CDA1452825). J.X. is supported by the grant from the Science and Technology Commission of Shanghai Municipality (23410750100). The proteomics experiment was performed at the Integrated Mass Spectrometry Shared Resource at the Beckman Research Institute, which was supported by the NIH (P30CA033572). The tissue histology analysis was performed at the Cell Biology and Morphology Core at the Boston-Area Diabetes and Endocrinology Research Center, which was supported by NIH (P30DK135043).

## Author contributions

Conceptualization: G.L., S.D., J.X., Methodology: T.H., J.L, G.L., S.D., J.X., Investigation: G.L., T.H., J.L., L.S., K.V.A., P.G., A.P., R.S., C.X, E.C., X.Y., R.X., Y.Z., Z.Y., Data analysis: G.L., K.V.A., A.C.M., T.H., K.G., S.L., P.P., Funding acquisition: S.D., G.L., Supervision: G.L., S.D., J.X., Writing – original draft: G.L., T. H., S.D., Writing – review & editing: all authors.

## STAR METHODS

### Isolation and culture of primary cardiac fibroblasts and cardiomyocytes

Primary neonatal rat ventricular cardiomyocytes (NRVMs) and cardiac fibroblasts (NRCFs) were isolated from 1–2-day-old Sprague-Dawley rat pups using serial enzymatic digestion as previously described^55^. Briefly, neonatal hearts were excised under sterile conditions, and atria were removed. Ventricular tissues were washed in adsorption solution (ADS) and minced into small fragments. The tissue was subjected to sequential enzymatic digestion in ADS containing collagenase (355 U/mg, 2.16 mg/mL) and pancreatin (3.24 mg/mL) at 37°C with gentle agitation (110-120 rpm). Digestion was performed through multiple sequential incubations, and the released cells from each digestion were collected into fetal bovine serum (FBS) to terminate enzymatic activity. Cell suspensions from all digestion rounds were pooled, centrifuged (1,200 rpm, 4 min), and resuspended in cardiomyocyte culture medium consisting of DMEM supplemented with 10% horse serum, 5% FBS, 1% penicillin/streptomycin, and 1% L-glutamine. The cell suspension was then subjected to differential pre-plating on uncoated culture dishes for approximately 60 min at 37°C to allow preferential attachment of cardiac fibroblasts while cardiomyocytes remained in suspension. The non-adherent cell fraction was collected and further purified by discontinuous Percoll density gradient centrifugation. Cells were layered onto a two-step Percoll gradient and centrifuged at 3,000 rpm for 30 min with low acceleration and deceleration settings. Purified cardiomyocytes were recovered from the medium fraction, washed with culture medium, centrifuged at 1,200 rpm for 4 min, resuspended in fresh cardiomyocyte culture medium, counted using a hemocytometer with trypan blue exclusion, and seeded onto culture-treated dishes for subsequent experiments. The adherent cells obtained during the pre-plating step were maintained as primary cardiac fibroblasts. After removal of non-adherent cardiomyocytes, attached fibroblasts were washed and cultured in Fibroblast Growth Medium 3 (PromoCell, C-23025) for subsequent experiments.

Primary adult rat cardiac fibroblasts (ARCFs) and adult human cardiac fibroblasts (hCFs) were obtained from Cell Applications (catalog no. R306-05a) and ScienceCell (catalog no. 6300), respectively. Both cells were cultured in Fibroblast Growth Medium 3 (PromoCell, C-23025).

NRVMs, NRCFs, ARCFs, and hCFs were seeded in either 24-well plates for immunofluorescence staining or 6-well plates for RNA isolation and allowed to attach overnight. The following day, cells were transfected with tDR mimics or control mimics using Lipofectamine RNAiMAX (Thermo Fisher, 13778150) according to the manufacturer’s instructions. After approximately 16 h, the transfection medium was replaced with fresh complete medium containing either 200 μM phenylephrine (PE) (Sigma-Aldrich, P6126) to induce cardiomyocyte hypertrophy or 10 ng/mL recombinant human TGF-β (Proteintech, H2824) to stimulate CF activation. Fresh PE- or TGF-β-containing medium was replenished every 24 h, and cells were stimulated for a total of 48 h.

### Human myocardial tissue sampling

This study complies with all relevant ethical regulations. All subjects provided written informed consent, and the Institutional Review Boards at Vanderbilt University Medical Center and Massachusetts General Hospital approved this study. Myocardial tissue was obtained as previously described ^60^. Briefly, following informed consent, left ventricular tissue was sampled from the basal septum of individuals undergoing heart transplant or left ventricular assist device implant (N=15) or from donor hearts unused for transplant (N=9), immediately snap frozen in liquid nitrogen, and stored long-term in −80⁰C. Patient information is described in **Table S4**.

### In vitro transfection of nucleic acid oligos

For the transfection of nucleic acid oligos, 50 nM tDR mimics (synthesized by Integrated DNA Technologies), 50 nM siRNAs (dsiRNAs, synthesized by Integrated DNA Technologies), 50 nM tDR biogenesis reporter (synthesized by Dharmacon), or 100 nM antisense oligonucleotides (ASOs, synthesized by Qiagen) were transfected into the cultured cells using Lipofectamine RNAiMax (Thermo Fisher, 13778150) by following the manufacturer’s instructions, unless otherwise specified. tDR mimics were synthesized at Integrated DNA Technologies in default mode, ending with 5′-OH and 3′-OH, with no additional modifications. The sequences of tDR mimics, ASOs, and siRNAs are listed in **Table S5.**

### Human induced pluripotent stem cell (hiPSC) culture, cardiomyocyte differentiation, and cardiac organoid generation

Human induced pluripotent stem cells (hiPSCs; iPS(IMR90) clone #4, WiCell Research Institute) were maintained on Matrigel-coated culture plates in mTeSR Plus medium (Stemcell Technologies), with daily medium changes. Cells were passaged using ReLeSR when they reached approximately 85% confluency and expanded until cardiac differentiation.

Directed differentiation of hiPSC-derived cardiomyocytes (hiPSC-CMs) was performed using a chemically defined Wnt modulation protocol optimized in our laboratory. Briefly, at ∼85% confluency (day 0), hiPSCs were washed with DPBS and cultured in RPMI 1640 supplemented with B27 minus insulin (RPMI minus) containing 6-8 μM CHIR99021. On days 1 and 2, fresh RPMI minus medium was added without replacing the existing medium. On day 3, cells were washed with DPBS and cultured in fresh RPMI minus supplemented with 3-4 μM IWR-1. Medium was subsequently replaced every 48 h. Beginning on day 9, cells were maintained in RPMI 1640 supplemented with B27 and 1% penicillin-streptomycin (RPMI Plus), with medium changes every 48 h. To enrich the cardiomyocyte population, cells underwent metabolic purification starting on day 15 by culturing in glucose-free RPMI supplemented with B27 and 1% penicillin-streptomycin for 6 days, with medium refreshed every 72 h. On day 22, differentiated hiPSC-CMs were dissociated and replated onto Matrigel-coated plates to remove cellular debris and establish confluent cardiomyocyte monolayers. Cells were maintained in RPMI Plus until further experiments. For tDR manipulation studies, hiPSC-CMs were replated into 6-well plates at approximately 80% confluency and transfected with tDR or control mimics using Lipofectamine RNAiMAX the following day. Cells were then incubated for 24 h before harvest or subsequent cardiac organoid fabrication.

Cardiac organoids were generated using agarose microwell culture as previously described ^61,62^. Briefly, 2% agarose microwells were cast using silicone molds and equilibrated with hiPSC-CM maintenance medium for at least 1 h before cell seeding. Dissociated hiPSC-CMs were resuspended at 2.33 × 10^6^ cells/mL, and 75 μL of the suspension was seeded into each agarose chamber, resulting in approximately 5 × 10^3^ cells per organoid. Cells were allowed to settle within the microwells for 1 h before adding RPMI Plus medium and were cultured for an additional 24 h to allow spontaneous aggregation into cardiac organoids. Cardiac hypertrophy was induced by treating organoids with endothelin-1 (ET-1; Sigma, E7764). One day after organoid formation, the culture medium was replaced with RPMI Plus containing 100 μM ET-1 or vehicle control. Fresh treatment medium was supplied after 24 h, and hypertrophic stimulation was maintained for a total of 48 h. At the endpoint, organoids were washed with DPBS, collected in TRIzol reagent, flash frozen in liquid nitrogen, and stored for subsequent RNA isolation and qRT-PCR analysis.

### Total RNA-seq

Total RNA-seq was performed as previously described^30^. Briefly, cells were lysed using TRIzol™ (Thermo Fisher, A33251) and then total RNAs were isolated using RNA Clean & Concentrator kit (Zymo Research, R1014) by following the manuals. cDNA libraries were constructed using the SMARTer Stranded Total RNA-Seq Kit v2 Pico Input Mammalian (Takara Bio, 634411) and sequenced using the Illumina NextSeq 2000 platform using a resolution of 100 base pairs with paired-end reads. Reads were trimmed to remove adapter sequences using Cutadapt (v2.10) and aligned to the Gencode mRatBN7.2 (rn7) genome using STAR (v2.7.8a). Gencode v38 gene annotations were provided to STAR to improve the accuracy of mapping. Quality control on both raw reads and adaptor-trimmed reads was performed using FastQC (v0.11.9) (www.bioinformatics.babraham.ac.uk/projects/fastqc), featureCounts (v2.0.2) was used to count the number of mapped reads to each gene. Heatmap3 was used for cluster analysis and visualization. Significantly differentially expressed genes with absolute fold change >=1.5 and FDR-adjusted p-value <= 0.05 were detected by DESeq2 (v1.30.1).

### Characterization of tDR-bound proteins and RNAs

To identify the binding proteins of tRNA-Asp-GTC-3’tDR, 50 nM Biotin-conjugated tRNA-Asp-GTC-3’tDR or control mimics were transfected into NRCFs or NRVMs using Lipofectamine RNAiMAX. 24 hours after transfection, the cells were pelleted after washing and proceeded with RNA pull-down using the Pierce™ Magnetic RNA-Protein Pull-Down Kit (Thermo Fisher, 20164). The enriched tDR binding partners were subjected to protein or RNA analyses.

Binding proteins were directly eluted using 1x Laemmli sample buffer (BioRad, 161-0747) and subjected to in-solution protein digestion using the SP3 protocol. The resulting tryptic peptides were directly loaded on a 50 cm C18 EasySpray column and separated by acidic reverse phase chromatography over a 75 min gradient. Data acquisition was carried out on a Thermo Orbitrap Eclipse mass spectrometer equipped with a FAIMS Pro device using the following settings: FAIMS CV at −40V, −60V, and −80V, MS1 resolution of 120K, full scan range of 375-1575 m/z, precursor isolation window of 1.6, MS2 scans in the iontrap from HCD fragmentation of precursor ions with 32% NCE and dynamic exclusion setting of 60 seconds. Data were acquired in top-speed mode with a cycle time of 1 sec per FAIMS CV. Data were searched in Proteome Discoverer v2.4 running Mascot v2.6 against a UniProtKB/Swiss-Prot rat protein sequence database. Search parameters included: Cys carbamidomethylation as a fixed and Met oxidation as a dynamic modification; precursor mass tolerance of 10 ppm and a fragment mass tolerance of 0.6 Da. PSM and peptide identifications were filtered to 1% FDR using the Percolator node in Proteome Discoverer v2.4.

Binding RNAs were collected using TriZol and isolated using RNA Clean & Concentrator kit (Zymo Research, R1014) by following the manuals. Isolated RNAs were then amplified using the SMARTer Stranded Total RNA-Seq Kit v2 Pico Input Mammalian (Takara Bio, 634411) and analyzed using qRT-PCR.

### Animal studies

Wild-type C57BL/6J mice (strain 000664) were purchased from The Jackson Laboratory and maintained under specific pathogen-free conditions. All animal procedures were approved by the Massachusetts General Hospital Institutional Animal Care and Use Committee and performed in accordance with the Guide for the Care and Use of Laboratory Animals (NIH Publication No. 85-23, revised 1996). Mice aged 8-14 weeks were used for all experiments. Group sizes were 3-4 mice for expression analyses and 5-11 mice per group for in vivo studies, as indicated in the figure legends. Myocardial infarction (MI), cardiac ischemia/reperfusion (I/R), and transverse aortic constriction (TAC) surgeries were performed as previously described with minor modifications^55^. Briefly, mice were anesthetized with 1–2% isoflurane in oxygen, intubated using a 22-GA Angiocath, and mechanically ventilated (SAR-830/AP Ventilator, USA) throughout the procedure. Animals were maintained on a 37°C heating pad, and the surgical site was depilated and sterilized with 10% Betadine followed by 70% ethanol. For MI and I/R surgeries, a left thoracotomy was performed at the third intercostal space to expose the heart. The left anterior descending (LAD) coronary artery was identified and ligated with a 6-0 silk suture approximately 3 mm distal to its origin. For the MI model, the ligature was permanently tied after confirmation of myocardial blanching distal to the ligation site. For the I/R model, the LAD was occluded using a slip knot for 45 min, followed by release of the ligature to allow reperfusion. For TAC surgery, a lateral thoracotomy was performed by bluntly separating the pectoral muscles with forceps and opening the first and second intercostal space. The transverse aorta was then carefully dissected free from the surrounding connective tissue and constricted by tying a 7-0 silk Prolene suture around a blunted 27-G needle, which was subsequently removed to achieve standardized constriction. Following each surgery, the chest cavity and skin incision were closed using 6-0 silk sutures. Mice were maintained on a warming pad until recovery from anesthesia, extubated after spontaneous respiration resumed, and monitored until fully ambulatory before being returned to their cages.

### In vivo administration of ASOs and tDR mimics

ASOs with fully phosphorothioate-modified backbones were synthesized by Qiagen. Modified ASOs targeting tRNA-Asp-GTC-3′ tDR or a negative control ASO were administered by intravenous injection (30 mg/kg body weight) every other day for 1 week.

Mimics of tRNA-Asp-GTC-3′ tDR and control were synthesized by Integrated DNA Technologies using standard parameters, yielding molecules with 5′-OH and 3′-OH termini without additional modifications. For in vivo delivery, PEI-based and lipid nanoparticle-based approaches were performed as previously described^30^. PEI-based delivery maintains a PEI nitrogen-to-RNA phosphate molar ratio of 8. For a 30 g mouse receiving 3 mg/kg body weight: tDR mimics (9 μl of 10 mg/ml stock in Nuclease-free water) were diluted in 91 μl of RNase-free 5% glucose solution in one tube, and 14.4 μl of 150 mM PEI-MAX (Polysciences, 24765) was diluted in 85.6 μl of RNase-free 5% glucose solution in a separate tube. The contents of both tubes were mixed thoroughly and incubated at room temperature for 15 min. The resulting 200 μl mixture was administered through intravenous injection. In the MI model, control mimics and Asp-GTC-3′tDR were administered one day before MI surgery, followed by twice-weekly dosing until 4 weeks post-surgery. In the cardiac I/R model, Asp-GTC-3′tDR or control mimics were administered immediately after reperfusion, followed by two additional injections during the first week after surgery. In the TAC model, Asp-GTC-3′tDR administration was initiated two weeks after surgery and maintained three times per week until week 10.

### Cardiac function assessment using serial echocardiography

Transthoracic echocardiography was performed with the Vevo 3100 Imaging System (Fuji Film Visual Sonics Inc., Canada) equipped with an MS-250 imaging transducer. The baseline cardiac function of mice was measured 3 days before surgery. Monitoring was initiated on postoperative day 1 and conducted on a weekly basis. The observation period was terminated at 4 weeks for MI, 8 weeks for I/R, and 12 weeks for TAC models. Mice were slightly anesthetized in a box with isoflurane. Their limbs were fixed in a supine position on the echo mat, and the chest hair was removed by depilating cream. Then, mice were anesthetized by inhalation of isoflurane (0.5 to 1%) mixed with oxygen to maintain the heart rate at approximately 500 to 600, and M-mode echocardiography was performed. The left ventricular internal diameter at end diastole (LV-IDd) and systole (LV-IDs) were obtained by measuring the long axis and the short axis. Accordingly, the cardiac parameters LV-EF, LV-FS, LV-EDV, and LV end-systolic volume (LV-ESV) were determined. The echocardiography measurement was carried out in a double-blind manner.

### Immunoblotting

Immunoblotting was performed as previously described ^30^. Briefly, cell pellets were resuspended in RIPA buffer (Thermo Fisher, 89900) containing Protease and Phosphatase Inhibitor Cocktail (Thermo Fisher, 78441), and PMSF, and lysed by sonication. Tissue samples were cut into small pieces and homogenized in RIPA buffer containing Protease and Phosphatase Inhibitor Cocktail and PMSF, followed by sonication. The protein concentration of clarified cell or tissue lysate was then measured using the BCA protein assay kit (Thermo Fisher, 23227). 2 ∼ 10 μg of total protein per sample was separated by SDS-PAGE gel and transferred to a PVDF membrane. 5% BSA in TBST was used for blocking and primary antibody incubation. HRP-conjugated secondary antibodies were then applied, and the blots were developed using SuperSignal West Femto Maximum Sensitivity Chemiluminescent Substrate (Thermo Fisher, 34094). The antibodies used are listed in **Table S6**.

### Nuclear and cytoplasmic fractionation

Nuclear and cytoplasmic fractions were isolated from NRVMs and NRCFs using a simplified nuclear and cytoplasmic protein fractionation protocol. Briefly, cells were washed twice with ice-cold PBS, collected, and lysed in cytoplasmic extraction buffer (20 mM Tris-HCl pH 7.4, 10 mM KCl, 2 mM MgCl_2_, 1 mM EGTA, 0.5 mM DTT) supplemented with protease inhibitors. Following centrifugation, the supernatant containing the cytoplasmic fraction was collected. The remaining nuclear pellet was washed and subsequently lysed in nuclear extraction buffer (20 mM Tris-HCl pH 7.4, 150 mM KCl, 2 mM MgCl_2_, 1 mM EGTA, 0.5 mM DTT) supplemented with protease inhibitors to obtain the nuclear fraction. An aliquot of the unfractionated cell lysate was retained as the input sample. Equal amounts of protein from the input, nuclear, and cytoplasmic fractions were analyzed using immunoblotting.

### Puromycin incorporation assay

Global protein synthesis rate was assessed using the puromycin incorporation assay. Cultured cells were treated with 10 μg/mL puromycin for 10 min at 37 ℃ before harvest. Cells were then washed with ice-cold PBS and lysed in RIPA buffer supplemented with protease inhibitors. Equal amounts of protein were subjected to SDS-PAGE and probed with an anti-puromycin antibody (Sigma-Aldrich, MABE343).

### Seahorse

The Seahorse XF Analyzer-based method for assessing cellular oxidative phosphorylation (OCR). Load pretreated cells (NRVM, 1.25×10^5^ cells/mL, NRCF, 0.625×10^5^ cells/mL) into the plate. Background correction wells (A1, A12, H1, H12) received 80 μL of medium without cells. The plate was left undisturbed in a tissue culture hood for 1 h to allow cells to settle uniformly, then transferred to a 37 °C, 5% CO₂ incubator for overnight attachment. Before the day of the assay, a Seahorse XFe96 probe plate was hydrated overnight at 37 °C in a non-CO₂ incubator with 200 μL of sterile water per well. On the assay day, the water was replaced with 200 μL of XF Calibrant (Agilent, 100840-000) per well and the probe plate was incubated for 60 min at 37 °C in a non-CO_2_ incubator. Assay medium (Agilent Seahorse XF DMEM Assay Medium, pH 7.4, 103575-100) was supplemented with 10 mM glucose, 1 mM pyruvate, and 2 mM glutamine, warmed to 37 °C. XF Cell Mito Stress Test Kit (Agilent, 103015-100): after removing 60 μL of culture medium, cells were gently washed twice with 200 μL of assay medium, yielding a final volume of 180 μL per well. The cell plate was then incubated at 37 °C in a non-CO₂ incubator for 60 min to degas. Meanwhile, drugs (oligomycin, FCCP, and rotenone/antimycin A, Rot/AA) were prepared according to the kit instructions, dissolved in assay medium, and loaded into the injection ports of the hydrated probe plate (port A: oligomycin, port B: FCCP, port C: Rot/AA). Data are exported via Seahorse Wave^®^ Software.

### Quantitative reverse transcription polymerase chain reaction

Quantitative reverse transcription polymerase chain reaction (qRT-PCR) was performed as previously described^63^. Briefly, cells or tissues were immediately lysed by adding TRIzol (Thermo Fisher, A33251), and total RNAs were isolated using the RNA Clean & Concentrator kit (Zymo Research, R1014 or R1018). 100 −1000 ng of total RNA was reverse transcribed to cDNA using the SuperScript III First-Strand Synthesis System (Thermo Fisher, 18080051) and then proceeded to qPCR using a QuantStudio 6 Flex Real-Time PCR System (Thermo Fisher) with SsoAdvanced Universal SYBR Green Supermix (BioRad, 1725275). The sequences of qPCR primers used are listed in **Table S7**.

### Northern blotting

Northern blotting was performed as previously reported ^30^. Briefly, denatured 0.1 ∼ 2 μg total RNAs were separated by 15% Criterion TBE-Urea PreCast Gels (Bio-Rad, 3450092 or 3450093). The gels were stained with SYBR-Gold (ThermoFisher, S11494) and transferred onto a positively charged Nylon membrane (Sigma-Aldrich, 11209299001) using the Trans-Blot Turbo Transfer System (Bio-Rad). The membranes were then crosslinked with EDC (Sigma-Aldrich, E7750) at 60°C for 1∼2 hours and prehybridized with ULTRAhyb Ultrasensitive Hybridization Buffer (Thermo Fisher, AM8669). 50 pmol/ml Biotin-labeled LNA-modified DNA probes (synthesized by Qiagen or Genscript) were used for hybridization at 37°C overnight. After washing sequentially with low-stringent buffer, high-stringent buffer, and 1× SSC, the blots were then processed and developed using the Chemiluminescent Nucleic Acid Detection Module Kit (Thermo Fisher, 89880). For all Northern blots, we used a single probe capable of detecting both full-length tRNAs and their corresponding tDRs within one intact blot.

### Histology and immunofluorescence staining

For paraffin embedding, hearts were fixed in 4% paraformaldehyde (PFA) overnight at 4°C, dehydrated through graded ethanol and xylene, embedded in paraffin, and sectioned at 5 μm. Paraffin sections were stained with the Masson’s Trichrome Stain Kit (Sigma-Aldrich, HT15) or Picrosirius Red Stain Kit (Abcam, ab150681) according to the manufacturers’ instructions to evaluate cardiac fibrosis. For immunofluorescence staining, paraffin sections were deparaffinized, rehydrated, and subjected to heat-mediated antigen retrieval in citrate buffer. After washing with PBS, sections were blocked with goat serum for 30 min at room temperature and incubated with primary antibodies overnight at 4°C. The following day, sections were washed with PBS and incubated with the appropriate fluorophore-conjugated secondary antibodies for 2 h at room temperature.

For frozen-section immunofluorescence, hearts were fixed in 4% PFA overnight, cryoprotected in 20% sucrose, embedded in optimal cutting temperature (OCT) compound, and sectioned at 5–10 μm. Frozen sections were permeabilized with 0.1% Triton X-100 in PBS for 10 min, blocked with 10% goat serum for 30 min at room temperature, and incubated with primary antibodies overnight at 4°C. After washing with PBS, sections were incubated with the appropriate fluorophore-conjugated secondary antibodies for 2 h at room temperature. Nuclei were counterstained and coverslips were mounted using ProLong™ Glass Antifade Mountant with NucBlue Stain (Thermo Fisher Scientific, P36981). Fluorescence images were acquired using a Leica SP8 confocal microscope.

For immunofluorescence staining of cultured cells, cells cultured in 24-well plates were fixed with 4% PFA for 15 min at room temperature, washed with PBS, permeabilized with 0.2% Triton X-100 in PBS for 10 min, and blocked with 10% goat serum in PBS for 1 h. Cells were then incubated with primary antibodies overnight at 4°C, followed by the appropriate fluorophore-conjugated secondary antibodies for 2 h at room temperature. Nuclei were counterstained with Hoechst 3342 (Thermo Fisher Scientific, H3570), and fluorescence images were acquired using a Leica SP8 confocal microscope.

### Statistical analysis

Quantitative densitometric analysis of immunoblotting and Northern blotting images was performed using BioRad ImageLab (version 6.1.0). Statistical analyses were performed using GraphPad Prism software (version 11). The data from immunoblotting, Northern blotting, and qRT-PCR are expressed as mean ± SEM, and their statistical significance was assessed using the unpaired two-tailed Student’s t test or the Mann-Whitney test, as annotated in the figure legend. Echocardiographic data are expressed as mean ± SEM, and their statistical significance was assessed using two-way ANOVA. For the transcriptomics data, differential expression analysis was performed using DESeq2 to generate Benjamini-Hochberg-corrected P-values (padj) to assess statistical significance. For proteomics data, protein abundances were normalized using variance stabilization normalization (vsn package) in R v3.5.2. Statistical significance in protein abundances between the control and tRNA-Asp-GTC-3′tDR pulldown experiment was assessed using an unpaired two-tailed Student’s t-test. Proteins with P value < 0.05 and log₂ fold-change (tRNA-Asp-GTC-3′tDR/control) > 1 were selected as significantly interacting proteins in the tRNA-Asp-GTC-3′tDR pulldown experiment. For patient and animal tissue analyses, the Mann-Whitney test was used to assess the statistical significance of the means between the two groups. The criterion for statistical significance was P < 0.05 (*P < 0.05, **P < 0.01, ***P < 0.001).

