## Supplement Information for "A tRNA-derived small RNA orchestrates cardioprotection through distinct cell-type-specific mechanisms"

### Supplemental information

**The PDF file includes:**

Figs. S1 to S8
Tables S1 to S7

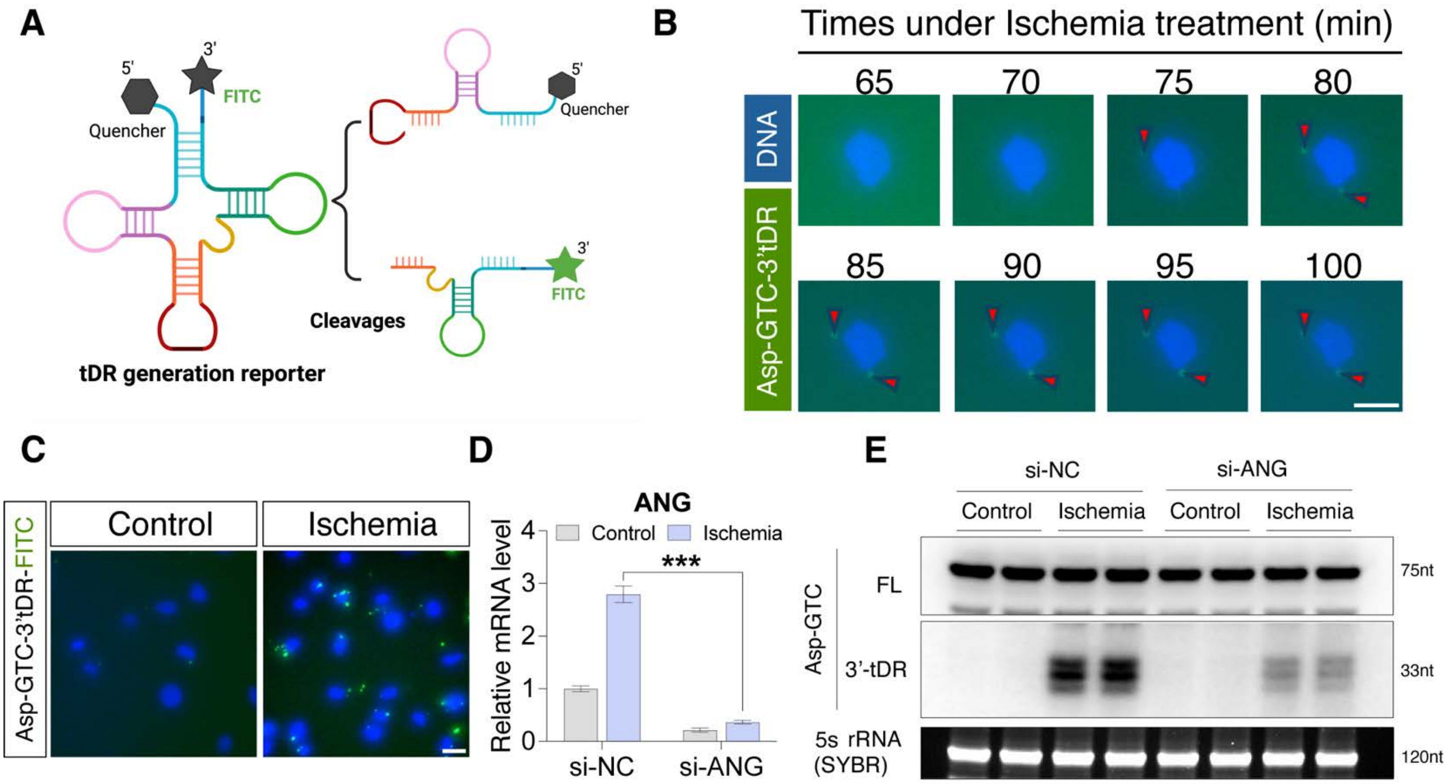


**Figure S1. Validation of Asp-GTC-3′tDR biogenesis in CFs under ischemic conditions.**

**(A)** Schematic illustration of the self-quenched Asp-GTC-3′tDR biogenesis reporter. **(B)** Representative live-cell fluorescence of NRCFs transfected with the Asp-GTC-3′tDR biogenesis reporter and subjected to ischemia for the indicated times. Scale bar, 50 μm. **(C)** Representative fluorescence of NRCFs transfected with the Asp-GTC-3′tDR biogenesis reporter under control or ischemic conditions for 6 hours. Scale bar, 100 μm. **(D)** qPCR analysis of ANG mRNA levels in NRCFs transfected with control siRNA (si-NC) or ANG-targeting siRNA (si-ANG) under control or ischemic conditions. **(E)** Northern blot analysis of Asp-GTC-3′tDR levels in NRCFs transfected with si-NC or si-ANG under control or ischemic conditions. Data are presented as mean ± SEM. Statistical significance was determined by the unpaired two-tailed Student's t-test. ****P* < 0.001.

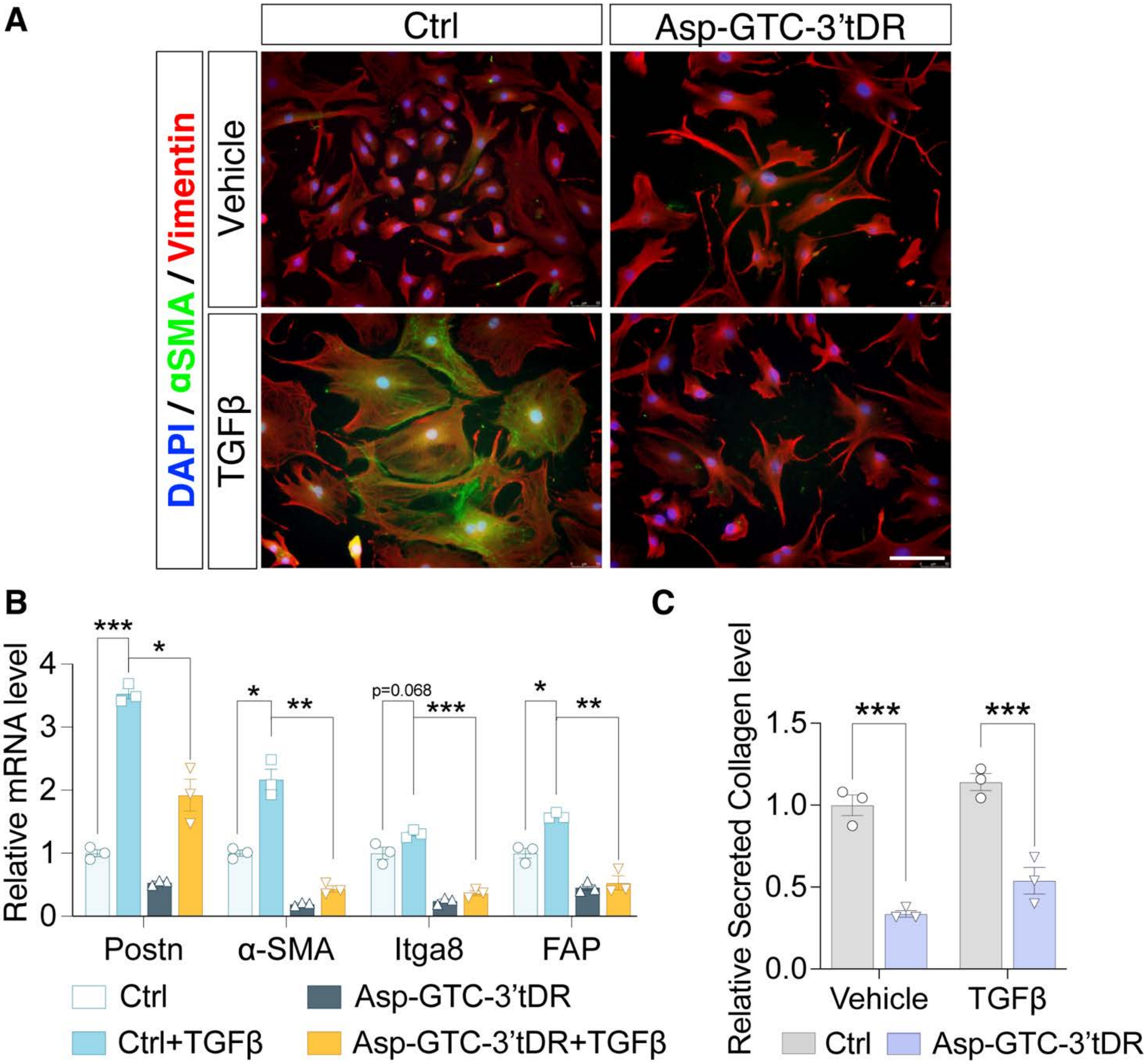


**Figure S2. Asp-GTC-3′tDR attenuates myofibroblast activation in adult rat CFs.**

**(A)** Immunofluorescence staining of adult rat CFs transfected with Ctrl or Asp-GTC-3′tDR mimic and treated with vehicle or TGFβ for 48 hours. Cells were stained for α-SMA (green), Vimentin (red), and nuclei (DAPI, blue). Scale bar, 100 μm. **(B)** qPCR analysis of myofibroblast activation-associated gene expression in adult rat CFs transfected with Ctrl or Asp-GTC-3′tDR mimic and treated with vehicle or TGFβ for 48 hours. **(C)** Secreted collagen levels in culture supernatants of adult rat CFs transfected with Ctrl or Asp-GTC-3'tDR mimics and treated with vehicle or TGFβ for 48 hours. Data are presented as mean ± SEM. Statistical significance was determined by the unpaired two-tailed Student's t-test. *$P < 0.05$, ** $P < 0.01$, ***$P < 0.001$.

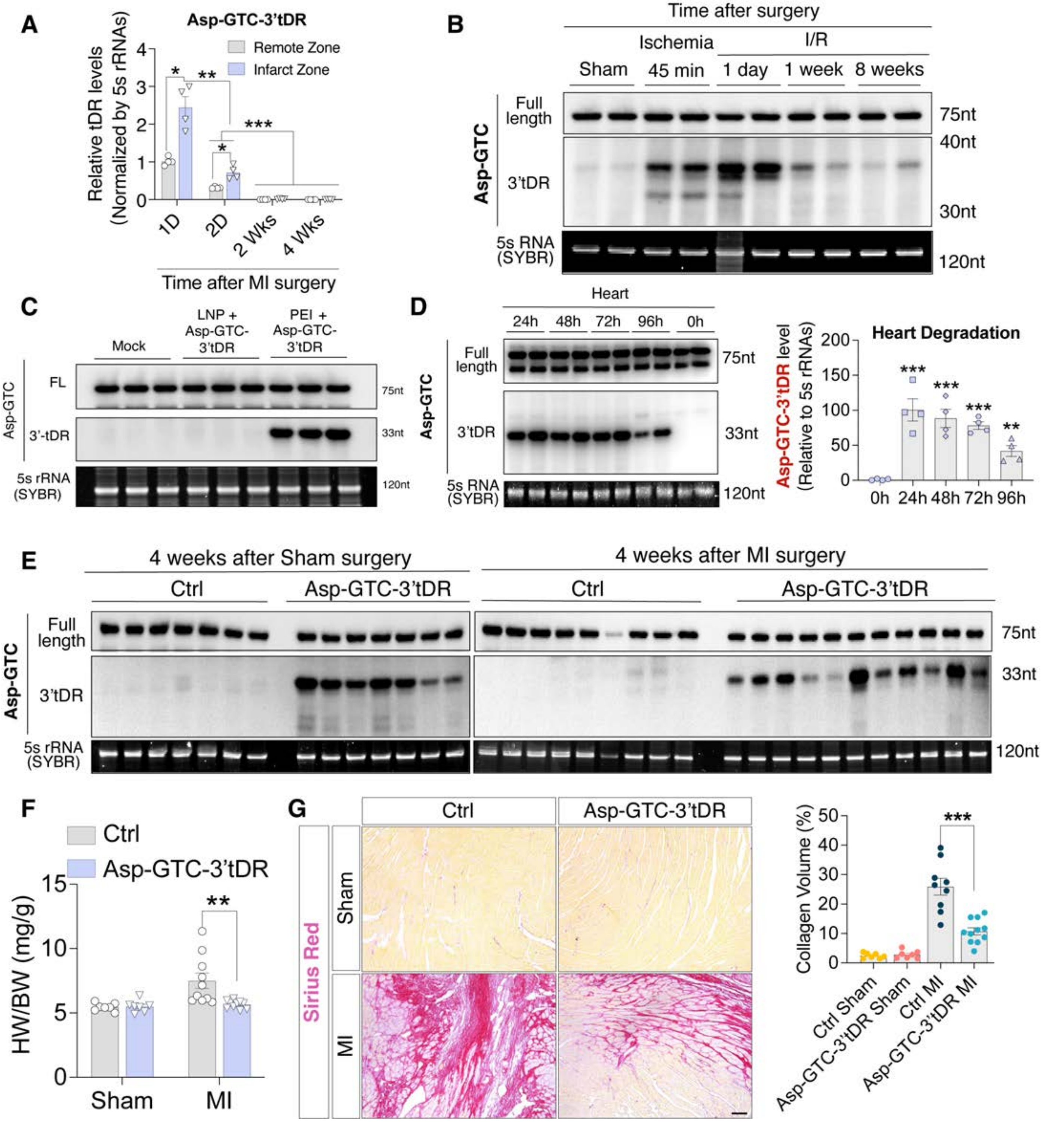


**Figure S3. Asp-GTC-3′tDR is induced by cardiac ischemia and attenuates post-MI cardiac remodeling in vivo.**

**(A)** Quantification of Asp-GTC-3′tDR levels in remote and infarct zones at the indicated time points following MI surgery. **(B)** Northern blot analysis of Asp-GTC-3′tDR levels in mouse hearts at the indicated time points following cardiac I/R injury. **(C)** Northern blot analysis comparing cardiac delivery of synthetic Asp-GTC-3′tDR by lipid nanoparticle (LNP)-based DharmaFECT versus polymer nanoparticle-based PEI. **(D)** Northern blot analysis and quantification of Asp-GTC-3′tDR mimic persistence in mouse hearts after a single administration of PEI-formulated Asp-GTC-3′tDR mimics. **(E)** Northern blot analysis of cardiac Asp-GTC-3′tDR levels in sham- and MI-operated mice treated with Ctrl or Asp-GTC-3′tDR mimics at 4 weeks. **(F)** Heart weight-to-body weight (HW/BW) ratios in sham and MI mice treated with control or Asp-GTC-3′tDR mimics. **(G)** Representative Sirius Red staining and quantification of collagen deposition in sham and MI hearts treated with Ctrl or Asp-GTC-3′tDR mimics. Scale bar, 100 μm. Data are presented as mean ± SEM. Statistical significance was determined by the unpaired two-tailed Student's t-test. $*P < 0.05$, $**P < 0.01$, $***P < 0.001$.

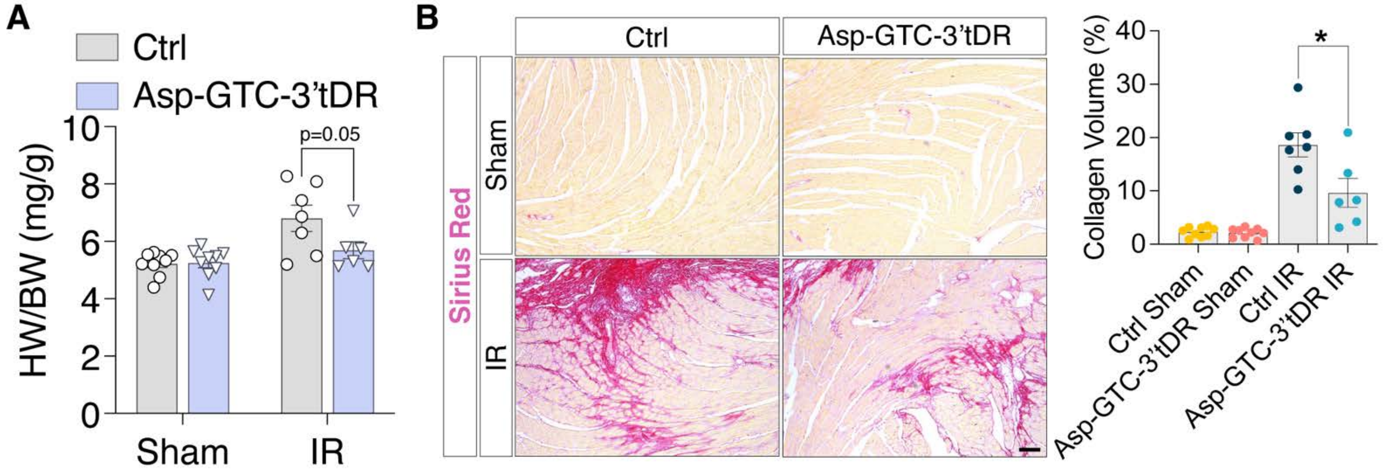


**Figure S4. Limited Asp-GTC-3′tDR administration attenuates cardiac fibrosis following cardiac I/R injury.** **(A)** Heart weight-to-body weight (HW/BW) ratios in sham and I/R mice treated with Ctrl or Asp-GTC-3′tDR mimics. **(B)** Representative Sirius Red staining and quantification of collagen deposition in sham and I/R hearts treated with Ctrl or Asp-GTC-3′tDR mimics. Scale bar, 100 μm. Data are presented as mean ± SEM. Statistical significance was determined by the unpaired two-tailed Student’s t-test. $^*P < 0.05$.

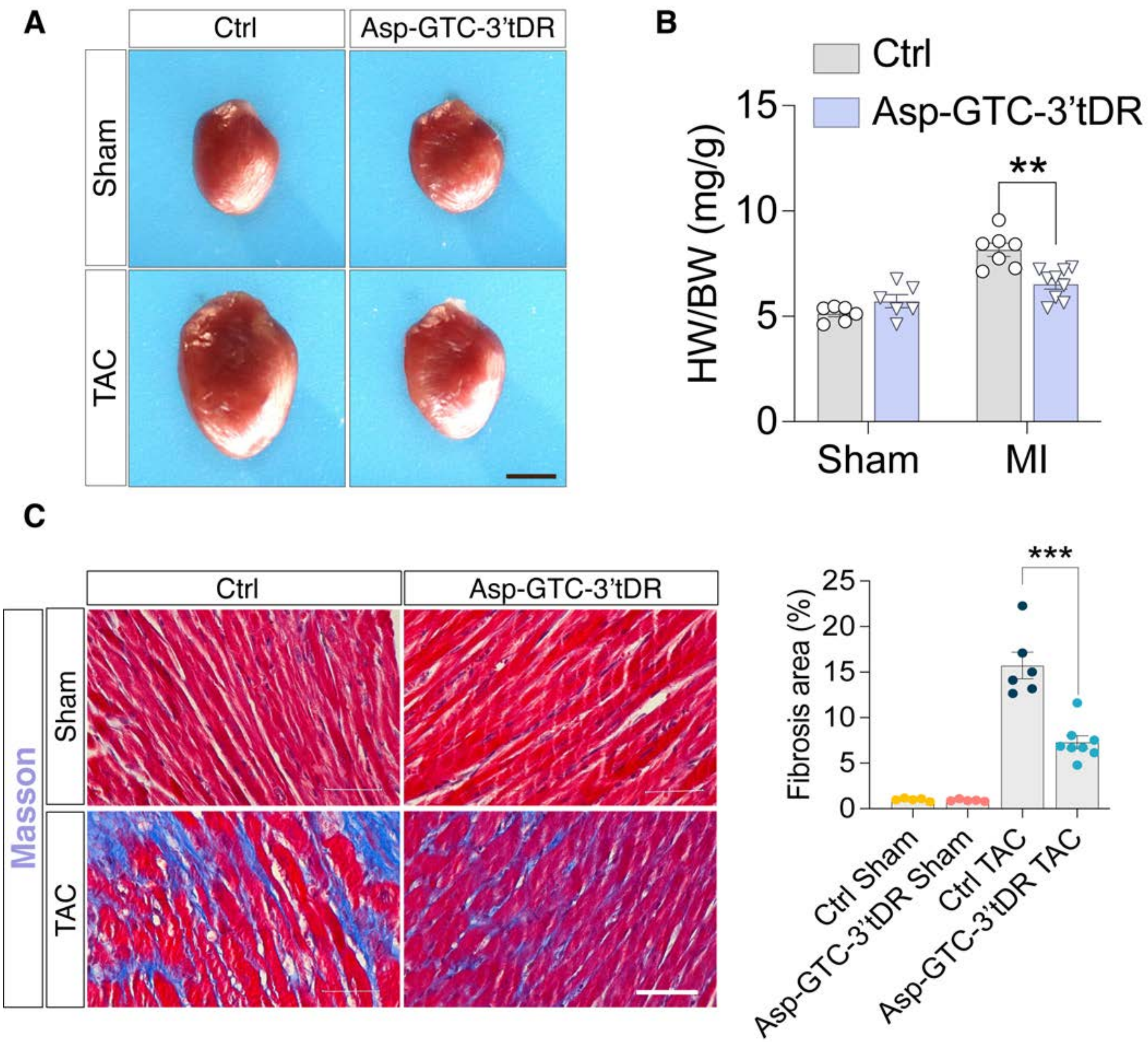


**Figure S5. Asp-GTC-3′tDR mitigates pathological cardiac remodeling in the murine TAC model.**
**(A)** Representative gross morphology of hearts harvested from sham and TAC mice treated with Ctrl or Asp-GTC-3′tDR mimics. Scale bar, 1 cm. **(B)** Heart weight-to-body weight (HW/BW) ratios in sham and TAC mice treated with Ctrl or Asp-GTC-3′tDR mimics. **(C)** Representative Masson's trichrome staining and quantification of fibrotic area in sham and TAC hearts treated with Ctrl or Asp-GTC-3′tDR mimics. Scale bar, 100 μm. Data are presented as mean ± SEM. Statistical significance was determined by the unpaired two-tailed Student's t-test. ** $P < 0.01$, *** $P < 0.001$.

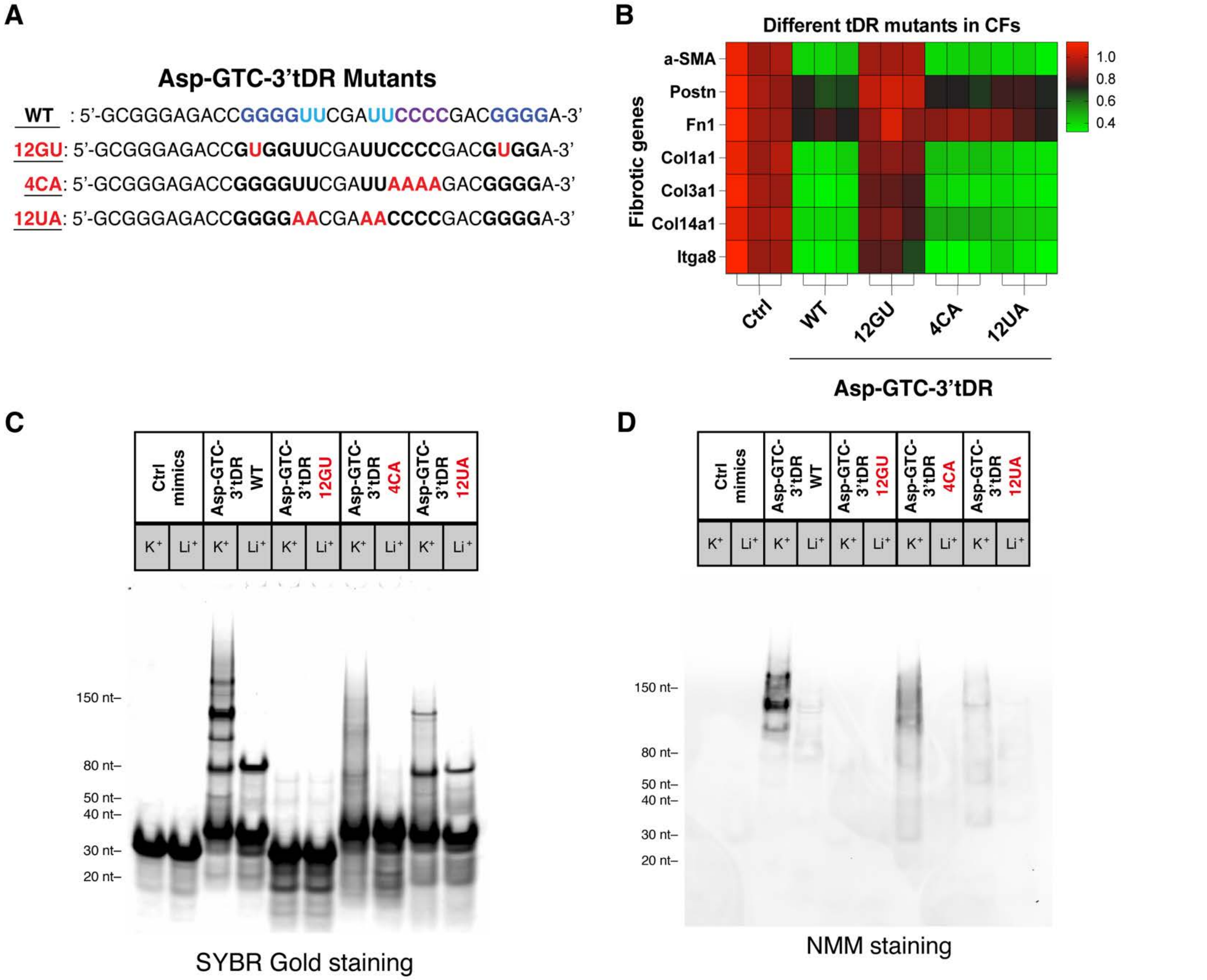


**Figure S6. G4 structure is required for the antifibrotic activity of Asp-GTC-3′tDR.**

**(A)** Sequences of wild-type (WT) Asp-GTC-3′tDR and the indicated mutants (12GU, 4CA, and 12UA). 12GU, converting the second guanine in each oligo-guanine motif to uridine; 4CA, replacing the 4 cytosines in the TΨC stem with adenines; and 12UA, substituting the four uridines in the TΨC loop with adenines. **(B)** Heatmap showing fibrosis-associated gene expression in NRCFs transfected with Ctrl, WT Asp-GTC-3′tDR, or the indicated Asp-GTC-3′tDR mutants. **(C)** Electrophoretic mobility shift analysis of intermolecular assemblies under G4 permissive (K+) or G4-nonpermissive (Li+) conditions for mimics of Ctrl, Asp-GTC-3′tDR, and its three mutants (12GU, 4CA, and 12UA). SYBR gold was used to stain the gels. **(D)** NMM staining of intermolecular assemblies under K+ or Li+ conditions for mimics of Ctrl, Asp-GTC-3′tDR, and its three mutants (12GU, 4CA, and 12UA).

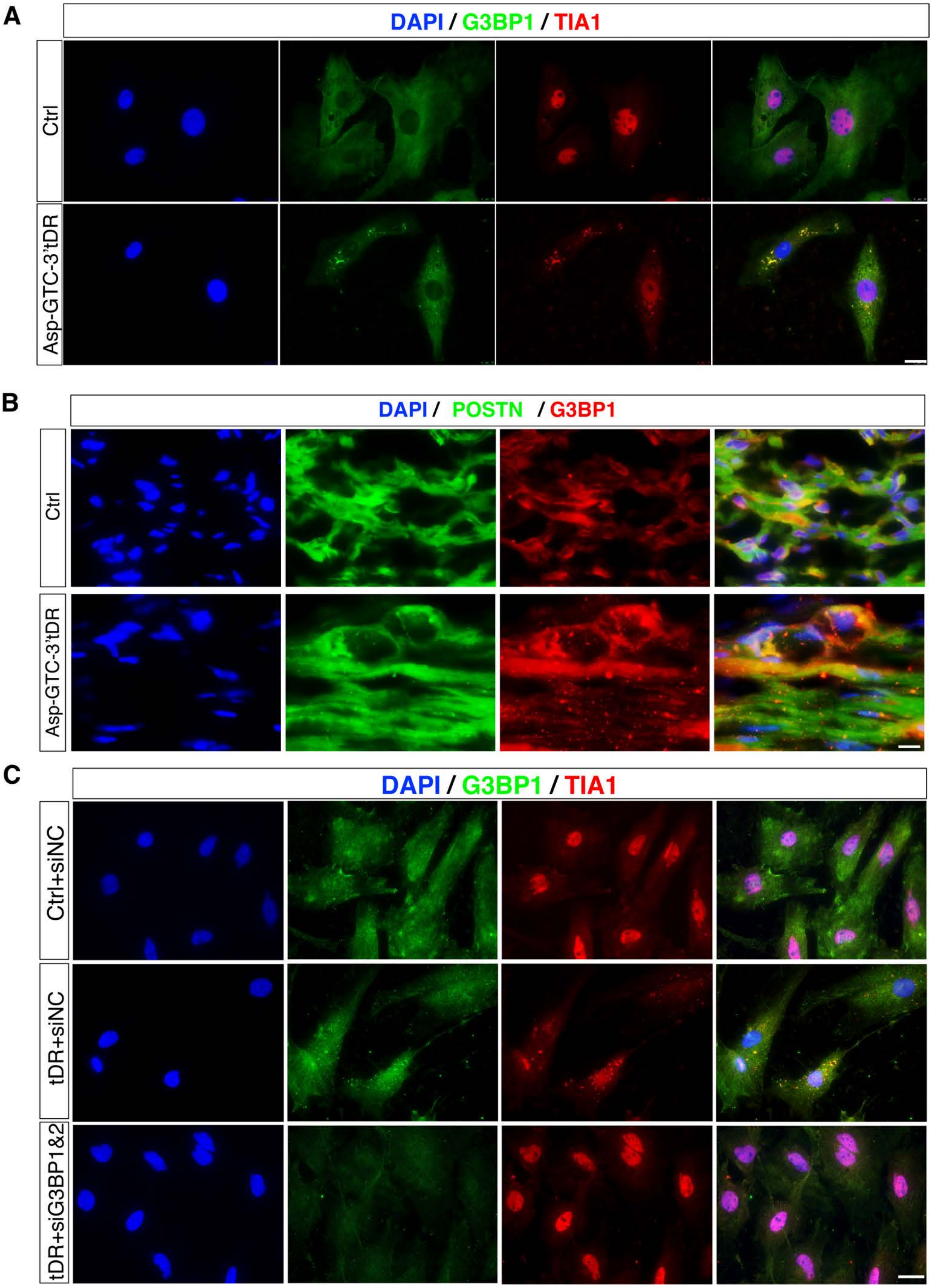


**Figure S7. Asp-GTC-3′tDR promotes SG assembly in CFs both in vitro and in vivo.**

(**A**) Representative immunofluorescence of G3BP1- and TIA1-positive SGs in adult rat CFs 24 hours after transfection with Ctrl or Asp-GTC-3′tDR mimics. Scale bar, 20 μm. (**B**) Representative immunofluorescence of POSTN and G3BP1 in the infarcted zone of mouse MI hearts treated with Ctrl or Asp-GTC-3′tDR mimics. Scale bar, 20 μm. (**C**) Representative immunofluorescence of G3BP1- and TIA1-positive SGs in NRCFs transfected with Ctrl or Asp-GTC-3′tDR mimic after co-silencing of G3BP1 and G3BP2 (siG3BP1&2). Scale bar, 20 μm.

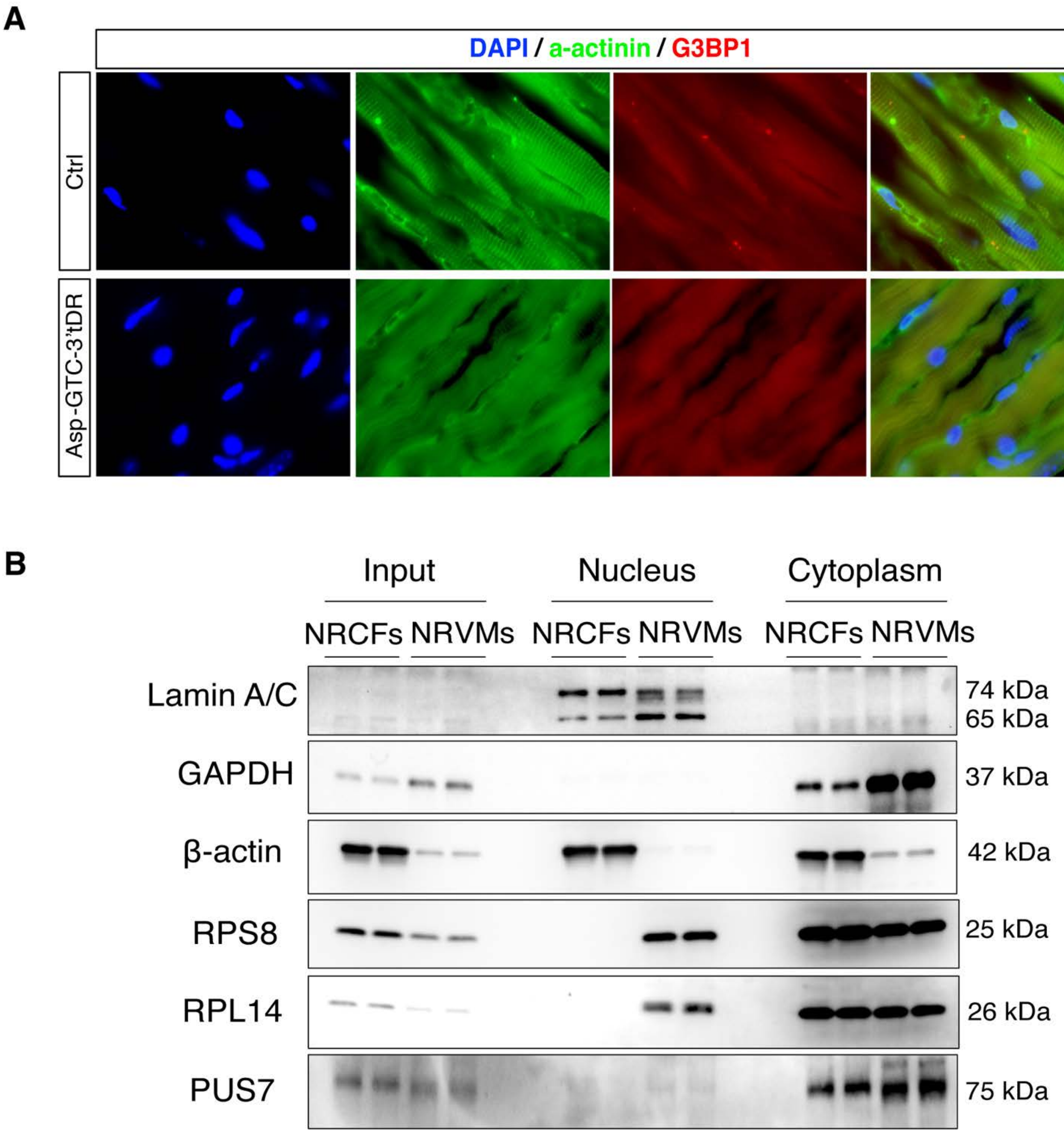


**Figure S8. Distinct regulatory pathways of Asp-GTC-3′tDR in CMs and CFs.**
(**A**) Representative immunofluorescence of α-actinin (green), G3BP1 (red), and DAPI (blue) in mouse MI hearts treated with Ctrl or Asp-GTC-3′tDR mimics, showing that Asp-GTC-3’tDR did not enhance SG formation in CMs. Scale bar, 20 μm. (**B**) Immunoblot analysis of subcellular abundance of ribosomal proteins, GAPDH, and PUS7, in total, nuclear, and cytoplasmic fractions from NRCFs and NRVMs. Lamin A/C and GAPDH served as nuclear and cytoplasmic fraction markers, respectively.

**Table S1. List of top 24 hypoxia-regulated tDRs in NRCFs.**

| Name | BaseMean | log2FoldChange | padj |
|---|---|---|---|
| tDR-1:32-Glu-TTC-4 | 2202.2035 | 3.7077632 | 1.04E-142 |
| tDR-1:32-Glu-CTC-1 | 774.943543 | 2.33459695 | 9.53E-133 |
| tDR-1:32-Asp-GTC-2-M2 | 1906.60743 | 2.67984207 | 1.48E-92 |
| tDR-39:72-Asp-GTC-2 | 625.498235 | 2.51153307 | 2.10E-65 |
| tDR-1:32-Gly-GCC-2-M2 | 887.750281 | 2.24010263 | 2.92E-59 |
| tDR-1:34-Glu-CTC-1 | 1056.41506 | 1.94849409 | 1.51E-50 |
| tDR-40:72-Asp-GTC-2 | 507.478498 | 3.65477253 | 1.43E-30 |
| tDR-2:32-Glu-TTC-4 | 439.576339 | 12.3483394 | 9.60E-24 |
| tDR-39:75-Glu-TTC-3-M2 | 381.586102 | 12.1438734 | 8.98E-23 |
| tDR-2:33-Glu-TTC-4 | 213.146453 | 11.3044736 | 5.64E-20 |
| tDR-39:74-Glu-TTC-3-M2 | 202.23727 | 11.2275974 | 1.09E-19 |
| tDR-2:32-Glu-TTC-2-M2 | 131.96493 | 10.6113372 | 1.32E-17 |
| tDR-1:32-Gly-CCC-1-M2 | 1332.99449 | 1.7955316 | 6.46E-17 |
| tDR-1:32-His-GTG-1 | 101.975809 | 10.2409084 | 1.98E-16 |
| tDR-2:31-Glu-TTC-4 | 68.2279366 | 9.66075091 | 3.04E-14 |
| tDR-39:74-Asp-GTC-2 | 66.7920432 | 9.62917336 | 8.51E-14 |
| tDR-1:32-Ser-GCT-1-M3 | 85.8898499 | 9.99223144 | 3.31E-13 |
| tDR-8:33-Glu-TTC-2-M3 | 45.1930704 | 9.06563405 | 5.55E-10 |
| tDR-1:33-Glu-TTC-4 | 6174.16061 | 2.24773313 | 1.78E-07 |
| tDR-1:33-Glu-TTC-2-M2 | 1865.92691 | 1.93748228 | 2.46E-06 |
| tDR-39:76-Ser-AGA-1 | 758.57108 | 1.27235308 | 2.46E-06 |
| tDR-1:31-Glu-TTC-4 | 597.99757 | 1.41454315 | 4.85E-06 |
| tDR-1:34-Glu-CTC-1-D5G | 370.061245 | 2.17453359 | 9.26E-06 |
| tDR-39:76-Ser-GCT-3 | 311.542706 | 1.55054116 | 0.00466966 |

Note: all tDRs were named by a standardized ontology, tDRnamer.

**Table S2. List of top 40 tRNA-Asp-GTC-3'tDR-binding proteins.**

| Accession ID | Gene | MW (kDa) | Log2FoldChange | pValue |
|---|---|---|---|---|
| G3V6S8 | Srsf6 | 39 | 4.676894586 | 2.89E-08 |
| Q5SGE0 | Lrpprc | 156.6 | 3.816037592 | 2.65E-06 |
| P62243 | Rps8 | 24.2 | 3.15453676 | 3.15E-06 |
| Q63507 | Rpl14 | 23.3 | 3.923538195 | 3.20E-06 |
| P13383 | Ncl | 77.1 | 2.375825839 | 4.46E-06 |
| P50878 | Rpl4 | 47.2 | 3.299795243 | 5.27E-06 |
| P05426 | Rpl7 | 30.3 | 3.113742507 | 5.91E-06 |
| P18484 | Ap2a2 | 104 | 3.077681611 | 6.31E-06 |
| D3ZBN0 | H1-5 | 22.6 | 3.52642821 | 6.87E-06 |
| P11250 | Rpl34 | 13.5 | 2.88421501 | 7.08E-06 |
| P23358 | Rpl12 | 17.8 | 2.77276531 | 1.00E-05 |
| O35821 | Mybbp1a | 152.2 | 3.990440231 | 1.05E-05 |
| P62752 | Rpl23a | 17.7 | 2.265915799 | 1.38E-05 |
| D3ZHV2 | Macf1 | 619.2 | 2.844429176 | 1.67E-05 |
| Q1RP77 | Nop16 | 21.1 | 2.252888314 | 1.97E-05 |
| Q6IMY8 | Hnrnpu | 87.7 | 2.38939527 | 3.08E-05 |
| P41123 | Rpl13 | 24.3 | 3.284247203 | 3.50E-05 |
| Q6AY09 | Hnrnph2 | 49.3 | 4.509220866 | 3.83E-05 |
| P17702 | Rpl28 | 15.8 | 6.001570556 | 4.39E-05 |
| P62919 | Rpl8 | 28 | 3.626837211 | 4.91E-05 |
| P12749 | Rpl26 | 17.3 | 2.890751594 | 4.95E-05 |
| P35427 | Rpl13a | 23.5 | 3.091582846 | 5.35E-05 |
| Q498T2 | Chtop | 26.5 | 1.789823739 | 6.68E-05 |
| P60123 | Ruvbl1 | 50.2 | 3.759508822 | 7.68E-05 |
| P21531 | Rpl3 | 46.1 | 4.221120591 | 7.81E-05 |
| P18395 | Csde1 | 88.8 | 3.321068227 | 7.95E-05 |
| Q9JIL3 | Ilf3 | 95.9 | 1.360686663 | 9.71E-05 |
| Q811S9 | Gnl3 | 60.6 | 3.796791243 | 1.10E-04 |
| Q66X93 | Snd1 | 101.9 | 2.151316633 | 1.25E-04 |
| P19945 | Rplp0 | 34.2 | 2.445919046 | 1.44E-04 |

| | | | | |
|---|---|---|---|---|
| O08837 | Cdc5l | 92.2 | 4.110547388 | 1.48E-04 |
| P61354 | Rpl27 | 15.8 | 2.534934165 | 1.91E-04 |
| P84092 | Ap2m1 | 49.6 | 2.363834294 | 2.16E-04 |
| Q62826 | Hnrnpm | 73.7 | 1.255306485 | 2.25E-04 |
| P29314 | Rps9 | 22.6 | 2.800913649 | 2.38E-04 |
| P63039 | Hspd1 | 60.9 | 1.115597331 | 2.73E-04 |
| P05765 | Rps21 | 9.1 | 1.546278751 | 3.05E-04 |
| Q9Z1H9 | Cavin3 | 27.9 | 4.349349152 | 3.26E-04 |
| P24049 | Rpl17 | 21.4 | 1.866202123 | 3.38E-04 |
| P61980 | Hnrnpk | 50.9 | 1.372346578 | 3.42E-04 |

**Table S3. The number of PUS7-targeting UGUAR motifs in tRNA-Asp-GTC-3'tDR regulated histone mRNAs.**

| Provided | Name | UNUAR | UGUAR |
|---|---|---|---|
| H1f2 | ENSRNOT00000147710 | 108 | 20 |
| H1f4 | ENSRNOT00000072564 | 7 | 1 |
| H1f5 | ENSRNOT00000151391 | 8 | 3 |
| H2ac4 | ENSRNOT00000110197 | 2 | 1 |
| H2ac12 | N/A | 1 | 1 |
| H2ax | ENSRNOT00000138971 | 10 | 3 |
| H2az2 | ENSRNOT00000058436 | 10 | 2 |
| H2bc18 | ENSRNOT00000095604 | 69 | 15 |
| H2bc4 | ENSRNOT00000097758 | 6 | 1 |
| H2bcl1 | ENSRNOT00000131854 | 87 | 12 |
| H3c13 | ENSRNOT00000110713 | 16 | 4 |
| H3c13 | ENSRNOT00000113668 | 10 | 3 |
| H4c1 | ENSRNOT00000147305 | 17 | 3 |
| H4c1 | ENSRNOT00000157059 | 2 | 1 |
| H4c8 | ENSRNOT00000080490 | 46 | 11 |
| H4c8 | ENSRNOT00000107328 | 30 | 9 |
| Hist1h2ail2 | ENSRNOT00000138608 | 21 | 3 |
| Hist1h2ao | ENSRNOT00000117583 | 20 | 5 |

**Table S4. Characteristics of patients with left ventricle tissue sampled for RNA analysis in Figure 7.**

| **Characteristic** | **Donor, N=9** | **ICM, N=15** |
|---|---|---|
| **Age at tissue procurement, years** | 48.2 ± 10.8 | 57.8 ± 8.8 |
| **Sex** | | |
| F | 5 (55.6%) | 1 (6.7%) |
| M | 4 (44.4%) | 14 (93.3%) |
| **Race** | | |
| White/Caucasian | 5 (55.6%) | 15 (100.0%) |
| not recorded/unknown | 4 (44.4%) | 0 (0.0%) |
| **Body mass index, $kg/m^2$** | — | 27.0 ± 5.0 |
| **Left ventricular ejection fraction** | | |
| <=40% (reduced) | 0 (0%) | 15 (100%) |
| 41-49% (mildly reduced) | 0 (0%) | 0 (0%) |
| >= 50% (preserved) | 9 (100%) | 0 (0%) |

*Mean ± SD; n (%)*

**Table S5. Sequences for tDR mimics, tDR probes, tDR ASOs, siRNAs.**

| Name | Applications | Species | Sequences |
|---|---|---|---|
| Ctrl mimics | RNA Mimics/GOF | N/A | AACGACAUACGCGUAUUAUACGC GAUUAACGAC |
| tRNA-Asp-GTC-5′tDR | RNA Mimics/GOF | Human, Mouse | UCCUCGUUAGUAUAGUGGUGAGU AUCCCCGCC |
| tRNA-Asp-GTC-3′tDR (33nt) | RNA Mimics/GOF | Human, Mouse | GCGGGAGACCGGGGUUCGAUUCC CCGACGGGGA |
| tRNA-Asp-GTC-3′tDR (12GU) | RNA Mimics/GOF | Human, Mouse | GCGGGAGACCGUGGUUCGAUUCC CCGACGUGGA |
| tRNA-Asp-GTC-3′tDR (4CA) | RNA Mimics/GOF | Human, Mouse | GCGGGAGACCGGGGUUCGAUUAA AAGACGGGGA |
| tRNA-Asp-GTC-3′tDR (12UA) | RNA Mimics/GOF | Human, Mouse | GCGGGAGACCGGGGAACGAAACC CCGACGGGGA |
| Probe for tRNA-Asp-GTC-5′tDR | DNA Probe/Northern blot | Human, Mouse | CTCACCACTATACTAACGAGGA |
| Probe for tRNA-Asp-GTC-3′tDR | DNA Probe/Northern blot | Human, Mouse | CCGTCGGGGAATCGAACC |
| Probe for tRNA-Glu-CTC-5′tDR | DNA Probe/Northern blot | Human, Mouse | CGAATCCTAACCACTAGACCAC |
| Probe for tRNA-Gly-GCC-5′tDR | DNA Probe/Northern blot | Human, Mouse | CTACCACTGAACCACCCATGC |
| Probe for tRNA-Glu-TTC-5′tDR | DNA Probe/Northern blot | Human, Mouse | CTAACCGCTAGACCATGTGGGA |
| ASO for tRNA-Asp-GTC-3′tDR | ASO/LOF | Human, Mouse | CGTCGGGGAATCGAACC |
| NC-ASO | ASO/LOF | | Qiagen catalog no. 339204-YCI0201821-FZA |
| tRNA-Asp-GTC tDR reporter | tDR biogenesis reporter | Human, Mouse | BHQ-1-UCCUCGUUAGUAUAGUGGUGAGU AUCCCCGCCUGUCACGCCGGGAG ACCGGGGUUCGAUUCCCCGACGG GGGAGCCA-FITC |
| DsiG3BP1-Seq1 | siRNA | Rat | GGAUGCUCACGCAACUCUGAAUGAC |
| DsiG3BP1-Seq2 | siRNA | Rat | GUCAUUCAGAGUUGCGUGAGCAUC CAC |
| DsiG3BP2-Seq1 | siRNA | Rat | UCAGUGAAUGUCAUACCAAAAUCCG |
| DsiG3BP2-Seq2 | siRNA | Rat | CGGAUUUUGGUAUGACAUUCACUGA AG |

**Table S6. Antibodies used in this study.**

| Antibody | Source | Catalog number | RRID number | Antibody type |
|---|---|---|---|---|
| α-SMA | Sigma | C6198 | AB_476856 | Mouse IgG2a |
| Vimentin | Abcam | Ab92547 | AB_10562134 | Rabbit IgG |
| cTNI | Abcam | Ab47003 | AB_869982 | Rabbit IgG |
| α-actinin | Abcam | Ab9465 | AB_307264 | Mouse IgG |
| Postn | Santa Cruze | Sc398631 | AB_2934053 | Mouse IgG |
| Connexin 43 | Abcam | Ab11370 | AB_297976 | Rabbit IgG |
| RPS8 | Abcam | Ab201454 | AB_2833046 | Rabbit IgG |
| RPL4 | Proteintech | 11302-1-AP | AB_2181909 | Rabbit IgG |
| RPL14 | Proteintech | 14991-1-AP | AB_2285319 | Rabbit IgG |
| NCL | Cell Signaling | 14574s | AB_2798519 | Rabbit IgG |
| LRPPRC | Proteintech | 21175-1-1AP | AB_10733879 | Rabbit IgG |
| Puromycin | Sigma | MABE343 | AB_2566826 | Mouse IgG2aκ |
| G3BP1 | Cell Signaling | 17798s | AB_2884888 | Rabbit IgG |
| TIA-1 | Invitrogen | MA5-26474 | AB_2725518 | Mouse IgG2b |
| UBAP2L | Cell Signaling | 40199T | AB_3754756 | Rabbit IgG |
| β-Actin | Sigma | A5316 | AB_476743 | Mouse IgG2a |
| LC3B | Cell Signaling | 3868 | AB_2137707 | Rabbit IgG |
| PUS1 | Thermofisher | PA5-66094 | AB_2665277 | Rabbit IgG |
| PUS7 | Thermofisher | PA5-54983 | AB_2646152 | Rabbit IgG |
| LC3A/B | Cell Signaling | 12741S | AB_2617131 | Rabbit IgG |
| GAPDH | Cell Signaling | 2118 | AB_561053 | Rabbit IgG |
| Lamin A/C | Cell Signaling | 2032 | AB_2136278 | Rabbit IgG |
| DHX36 | Novus Biologicals | NBP1-84286 | AB_11032598 | Rabbit IgG |
| Anti-Rabbit IgG/HRP | Agilent | P044801-2 | AB_2617138 | Goat IgG |
| Anti-Mouse IgG/HRP | Agilent | P044701-5 | AB_2617137 | Goat IgG |

| | | | | |
|---|---|---|---|---|
| Anti-Rat IgG (HRP) | Abcam | ab97057 | AB_10680316 | Goat IgG |
| Alexa Fluor® 488 Anti-Rabbit IgG | Thermofisher | A-21206 | AB_2535792 | Donkey IgG |
| Alexa Fluor® 594 Anti-Rabbit IgG | Thermofisher | A-21207 | AB_141637 | Donkey IgG |
| Alexa Fluor® 488 Anti-Mouse IgG | Thermofisher | A-21202 | AB_141607 | Donkey IgG |
| Alexa Fluor® 594 Anti-Mouse IgG | Thermofisher | A-21203 | AB_2535789 | Donkey IgG |

**Table S7. RT-PCR Primers used in this study.**

| Name | Species | Sequences |
|---|---|---|
| hmq-Actb-F | Human, Mouse, Rat | GACCTGTACGCCAACACAG |
| hmq-Actb-F | Human, Mouse, Rat | CTCAGGAGGAGCAATGATC |
| hq-18s-F | Human | GCCGCTAGAGGTGAAATTCT |
| hq-18s-F | Human | TCGGAACTACGACGGTATCT |
| rq-α-SMA-F | Rat | ACCATCGGGAATGAACGCTT |
| rq-α-SMA-R | Rat | CTGTCAGCA ATGCCTGGGTA |
| rq-Postn-F | Rat | CTCATAGTCGTATCAGGGGTCG |
| rq-Postn-R | Rat | ACACAGTCGTTTTCTGTCCAC |
| rq-Timp1-F | Rat | TGGCATCCTCTTGTTGCTATC |
| rq-Timp1-R | Rat | CCTTATAACCAGGTCCGAGTTG |
| rq-CTGF-F | Rat | GGGTCTCTTCTGCGACTTC |
| rq-CTGF-R | Rat | ATCCAGGCAAGTGCACTGGTA |
| rq-IL-11-F | Rat | AGCGCTGTCCTCTTGACCAG |
| rq-IL-11-R | Rat | GGAGTCCAGATTGTGGTCTCCATC |
| rq-TGFβ-F | Rat | TAATGGTGGACCGCAACAACG |
| rq-TGFβ-R | Rat | GGCACTGCTTCCCGAATGTCT |
| rq-Col1α1-F | Rat | AGCATGTCTGGTTTGGAGAG |
| rq-Col1α1-R | Rat | GTGATAGGTGATGTTCTGGGAG |
| rq-Col1α2-F | Rat | TGTTCGTGGTTCTCAGGGTAG |
| rq-Col1α2-R | Rat | TTGTCGTAGCAGGGTTCTTTC |
| rq-Col3α1-F | Rat | AGGGAACAACTGATGGTGCTACTG |
| rq-Col3α1-R | Rat | GGACTGCTGTGCCAAAATAAGAGA |
| rq-Col4α1-F | Rat | CCAGGAACGACTACTCCTACT |
| rq-Col4α1-R | Rat | CAA ACTGCACACCTGCTAATG |
| rq-Col8α1-F | Rat | CTCTACAGCTGCTGGGAATAC |
| rq-Col8α1-R | Rat | GTGGTATCTGAGGAGGGATTTG |
| rq-Timp1-F | Rat | TGTGCACAGTGTTTCCCTGT |
| rq-Timp1-R | Rat | TAGCCCTTCTCAGAGCCCAT |
| rq-MMP2-F | Rat | CTGTCTCCTGCTCTGTAGTTAATC |
| rq-MMP2-R | Rat | GATACGGTCAGCACCTTTCTT |
| rq-Itgα8-F | Rat | GGACCTGAACCAAGATGGATAC |
| rq-Itgα8-R | Rat | CTGGCGTTTCCGTTGTAAATG |
| rq-G3BP1-F | Rat | GTCTGCCTAGTGGGATTAAAGG |
| rq-G3BP1-R | Rat | ATGCTAAAGGACCTCAGAGAAAG |
| rq-G3BP2-F | Rat | GGCTTTCTTGTCAGTGGTTTG |
| rq-G3BP2-R | Rat | GCAGATGTTGCTGTGTGTAAAT |
| rq-H1F3-F | Rat | CGACGTGGAGAAGAACAACA |
| rq-H1F3-R | Rat | CGCCTTCTTGTTGAGTTTGAAG |
| rq-H1F4-F | Rat | CTCCGGCTCCTTCAAACTC |
| rq-H1F4-R | Rat | GTAGCCTTCTTGGGCTTCTT |
| rq-H2AC4-F | Rat | CCGCAAGGGCAACTACTC |

| | | |
|---|---|---|
| rq-H2AC4-R | Rat | GGATGATACGCGTCTTCTTGT |
| rq-H2AC12-F | Rat | CCGCAAGGGCAACTACTC |
| rq-H2AC12-R | Rat | ATGATGCGCGTCTTCTTGT |
| rq-H2AC20-F | Rat | CATCCGCAACGACGAAGA |
| rq-H2AC20-R | Rat | CTTAGCCTTGTGGCTCTCAG |
| rq-H2BC4-F | Rat | CTGTGAGGAAGAATGAGGGAAA |
| rq-H2BC4-R | Rat | CCAGGATTCCCAGAATGAAAGA |
| rq-H2BC18-F | Rat | AGACGTGATTGATCGGCTTTAC |
| rq-H2BC18-R | Rat | TAGACTCACGACCAGACCTTAC |
| rq-H4C1-F | Rat | AACATCCAGGGCATCACTAAG |
| rq- H4C1-R | Rat | AGGAACACCTTCAGCACAC |
| rq-H4CL1-F | Rat | GAGCATGCGAGGTAGCATATAA |
| rq-H4CL1-R | Rat | GAAAGCTGCATGGTTATGGATTAG |
| rq-H4C8-F | Rat | CCTGATTTCTGCCTAAGGGTTAT |
| rq-H4C8-R | Rat | TAAAGGCGGGATGGTGTAAAG |
| rq-H4C14-F | Rat | CGTGGTGTACTGAAGGTGTT |
| rq-H4C14-R | Rat | GCGCTTGAGAGCGTACA |
| rq-Itgα8-F | Rat | GGACCTGAACCAAGATGGATAC |
| rq-Itgα8-R | Rat | CTGGCGTTTCCGTTGTAAATG |
| mq-Postn-F | Mouse | AGGTGGCGATGGTCACTTAT |
| mq-Postn-R | Mouse | TGGCCTCTGGGTTTTCACTG |
| mq-α-SMA-F | Mouse | GTCCCAGACATCAGGGAGTAA |
| mq-α-SMA-R | Mouse | TCGGATACTTCAGCGTCAGGA |
| mq-FN1-F | Mouse | GTGTCTATGCTCTCAAGGA |
| mq-FN1-R | Mouse | CTAATAGTGATGGTGGTCTCT |
| mq-Col1α1-F | Mouse | CCGAGGTATGCTTGATCT |
| mq-Col1α1-R | Mouse | GACAGTCCAGTTCTTCATTG |
| mq-Col3α1-F | Mouse | TTCTTCTCACCCTTCTTCAT |
| mq-Col3α1-R | Mouse | GGCTCCTCATCACAGATTA |
| mq-Col8α1-F | Mouse | AAGAAGAGTCAAGCAAGGA |
| mq-Col8α1-R | Mouse | GTGGTGGTATCTGAGGAG |
| mq-MMP2-F | Mouse | GATTGACGCTGTGTATGAG |
| mq-MMP2-R | Mouse | CTTCTTGTTCTTACTCCAGTTAA |
| m/rq-FAP-F | Mouse, Rat | GATGGCAAGAGCAGAATATTTCAG |
| m/rq-FAP-R | Mouse, Rat | GTCAGAGTACCACATCGCC |
| m/r-ANP-F | Mouse, Rat | GAGCAAATCCCGTATACAGTGC |
| m/r-ANP-R | Mouse, Rat | ATCTTCTACCGGCATCTTCTCC |
| m/r-BNP-F | Mouse, Rat | GCTGCTGGAGCTGATAAGAGAA |
| m/r-BNP-R | Mouse, Rat | GTTCTTTTGTAGGGCCTTGGTC |
| m/r-α-MHC-F | Mouse, Rat | ACATCAGTCAGCAGAACA |
| m/r-α-MHC-R | Mouse, Rat | TTCCTCTAGCCTCTCACT |
| m/r-β-MHC-F | Mouse, Rat | GCTGTTATTGCTGCCATT |
| m/r-β-MHC-R | Mouse, Rat | TTATCATTCCGAACTGTC |

| | | |
|---|---|---|
| hq-Postn-F | Human | GAGCTTTACAACGGGCAAATAC |
| hq-Postn-R | Human | CTCCCTTGCTTACTCCCTTTC |
| hq-FN1-F | Human | CCACAGTGGAGTATGTGGTTAG |
| hq-FN1-R | Human | CAGTCCTTTAGGGCGATCAAT |
| hq-α-SMA-F | Human | GTGTTGCCCCTGAAGAGCAT |
| hq-α-SMA-R | Human | GCTGGGACATTGAAAGTCTCA |
| hq-CTGF-F | Human | AAA AGTGCATCCGTACTCCCA |
| hq-CTGF-R | Human | CCGTCGGTACATACTCCACAG |
| hq-ANP-F | Human | CAACGCAGACCTGATGGATTT |
| hq-ANP-R | Human | AGCCCCCGCTTCTTCATTC |
| hq-α-MHC-F | Human | GCCCTTTGACATTCGCACTG |
| hq-α-MHC-R | Human | GGTTTCAGCAATGACCTTGCC |
| hq-β-MHC-F | Human | CTTTGCTGTTATTGCAGCCATT |
| hq-β-MHC-R | Human | AGATGCCAACTTTCCTGTTGC |
| hq-H1-2-F | Human | AAGAAG AAGGCGGCCAAA |
| hq-H1-2-R | Human | GAAACTCCGCTACGCTCTTTA |
| hq-H2AC4-F | Human | CGCAATGACGAGGAGCTTAATA |
| hq-H2AC4-R | Human | TCACTTTCCCTTGGCCTTATG |
| hq-H2BC5-F | Human | CAGAAGAAGGACGGGAAGAAG |
| hq-H2BC5-R | Human | CCCATTGCCTTGGAAGAGAT |
| hq-H3C12-F | Human | CATCCGCAA ACTGCCATTTC |
| hq-H3C12-R | Human | CTTCAAAGAGACCCACCAGATAG |
| hq-H4C2-F | Human | GGCGTTCTCAAGGTGTTTCT |
| hq-H4C2-R | Human | TGAGCGCGTAAACCACATC |
| hq-H4C3-F | Human | GGATAACATCCAGGGCATTACA |
| hq-H4C3-R | Human | CACCTCGAGTCTCCTCATAGAT |
| hq-H1-4-F | Human | GAAGACTCCCGTGAAGAAGAAG |
| hq-H1-4-R | Human | TTGAGAGCGGCCAAAGATAC |
| hq-H2BC4-F | Human | GACACTGGCATCTCTTCCAA |
| hq-H2BC4-R | Human | GTCGAGCGCTTGTTGTAATG |
| hq-H2BC7-F | Human | CATGCCTGAACCTGCTAAGT |
| hq-H2BC7-R | Human | TACGCTTGCGCTTCTTACC |
| hq-H3C10-F | Human | TGATGGCGCTGCAAGAG |
| hq-H3C10-R | Human | GGATGTCCTTGGGCATGATAG |
| hq-CA9-F | Human | TCTACCTGGAGAGGAGGATCTA |
| hq-CA9-R | Human | CTTTGTCCCTGTGGGCATTAT |
| hq-BNIP3L-F | Human | ATGTCGTCCCACCTAGTCGAG |
| hq-BNIP3L-R | Human | TGAGGATGGTACGTGTTCCAG |
| hq-TRIB3-F | Human | CAAGGTTGAGGGACAGGATTAG |
| hq-TRIB3-R | Human | TAGAGTATGGACCTGGGATTGT |
